# Mcm2 promotes stem cell differentiation via its ability to bind H3-H4

**DOI:** 10.1101/2022.06.16.496513

**Authors:** Xiaowei Xu, Xu Hua, Kyle Brown, Xiaojun Ren, Zhiguo Zhang

## Abstract

Mcm2, a subunit of the Mcm2-7 helicase best known for its role in DNA replication, contains a histone binding motif that facilitates the transfer of parental histones following DNA replication. Here we show that Mcm2 is important for the differentiation of mouse embryonic stem (ES) cells. The Mcm2-2A mutation defective in histone binding impairs differentiation and disrupts the programmatic changes in gene expression and histone modifications during differentiation. Mcm2 localizes at transcription starting sites and the binding of Mcm2 at gene promoters is disrupted in both Mcm2-2A ES cells and neuro-precursor cells (NPCs). Reduced Mcm2 binding at bivalent chromatin domains containing repressive H3K27me3 and active H3K4me3 modifications in Mcm2-2A ES cells correlates with decreased chromatin accessibility at corresponding sites in NPCs. Together, our studies reveal a novel function of Mcm2 in ES cell differentiation, likely through manipulating chromatin landscapes at bivalent chromatin domains.

## Introduction

Embryonic stem (ES) cells are pluripotent cells that possess both the ability to self-renew and the potential to differentiate into lineage-specific cell types (De Los Angeles et al., 2015). The ability of ES cells to differentiate into specific lineage cell type both *in vivo* and *in vitro* opens exciting opportunities to study the events regulating the earliest stages of lineage specification during development (Keller, 2005). During mouse ES cell differentiation, pluripotency genes such as *Oct4*, *Sox2* and *Nanog* are silenced, whereas lineage-specific genes are up-regulated (Sha and Boyer, 2008). These dynamic changes in gene expression are regulated by transcription factors as well as chromatin factors (Young, 2011). For instance, the silencing of pluripotency genes (*Oct4*, *Sox2* and *Nanog*) is associated with both a dramatic reduction of tri-methylation of histone H3 lysine 4 (H3K4me3), a histone modification associated with active gene transcription, and an increase in H3K27me3, a repressive histone modification (Atlasi and Stunnenberg, 2017; Mikkelsen et al., 2007). In contrast, at the promoters of lineage specific genes H3K27me3 decreases while H3K4me3 increases (Bernstein et al., 2006; Harikumar and Meshorer, 2015). Although significant processes have been made to understand these dynamic changes in chromatin states during differentiation, the regulation of these dynamic changes remains underexplored.

Histone chaperones, a group of proteins best known for their roles in the assembly of DNA into nucleosomes following DNA replication and gene transcription, have been found to play multiple roles in stem cell maintenance and differentiation. For instance, chromatin assembly factor 1 (CAF-1), a histone chaperone involved in the deposition of newly synthesized H3-H4 onto replicating DNA, is important for cell fate maintenance (Ishiuchi et al., 2015; Smith and Stillman, 1989). Depletion of subunits of the CAF-1 complex in embryonic fibroblasts results in increased reprogramming efficiency into iPSC cells (Cheloufi et al., 2015). In ES cells, CAF-1 depletion leads to an increase in the percentage of 2C-like cells (2-cell-stage embryos), as well as defects in differentiation (Cheng et al., 2019; Ishiuchi *et al*., 2015). The role of CAF-1 in differentiation and the 2C-like state is linked to the function of CAF-1 in DNA replication-coupled nucleosome assembly. In contrast, Asf1a, a histone chaperone involved in the delivery of newly synthesized H3-H4 to CAF-1 in replication-coupled nucleosome assembly, as well as to HIRA, a histone chaperone involved in replication-independent (RI) nucleosome assembly (De Koning et al., 2007; English et al., 2006; Tagami et al., 2004), facilitates the expression of lineage-specific genes during ES cell differentiation, likely through regulating the disassembly of nucleosomes at bivalent chromatin domains (Gao et al., 2018). Moreover, histone H3.3 and its chaperone HIRA, facilitate the PRC2 silencing complex at developmental loci during differentiation (Banaszynski et al., 2013; Ray-Gallet et al., 2002). The H3.3-HIRA pathway also safeguards identities of differentiated cells, indicating its bimodal role in cell fate transition (Fang et al., 2018). Therefore, histone chaperones involved in deposition of newly synthesized H3-H4 play multiple roles in ES cell differentiation and maintenance.

In addition to histone chaperones involved in the deposition of newly synthesized histones, we and others have also uncovered a group of proteins that function in recycling parental histone H3-H4 following DNA replication. Pole3 and Pole4, two subunits of leading strand DNA polymerase ε, interact with H3-H4 and facilitate the transfer of parental histones to leading strands of DNA replication forks in both yeast and mouse ES cells (Li et al., 2020; Yu et al., 2018). On the other hand, Mcm2, a subunit of the Minichromosome Maintenance Proteins 2-7 (Mcm2-7) complex that plays an essential role in DNA replication as the replicative helicase (Tye, 1999), contains a histone binding domain (HBD) (Huang et al., 2015). Mutations at HBDs (Y81A and Y90A) of Mcm2 (Mcm2-2A) in both yeast and mouse ES cells leads to defects in the transfer of parental histones to lagging strands of DNA replication forks (Gan et al., 2018; Li et al., 2020; Petryk et al., 2018). Moreover, Mcm2 coordinates DNA replication progression and histone dynamics in complex with histone chaperone Asf1 (Groth et al., 2007). Besides its functions in DNA replication and histone deposition, Mcm2 also interacts with the carboxyl-terminal domain of RNA Pol II in *Xenopus* oocytes, and the Mcm2-7 complex is required for RNA Pol II-mediated transcription at some settings in mammalian cells (Snyder et al., 2009; Yankulov et al., 1999), indicating a possible role of Mcm2 in gene transcription.

Mouse ES cells with deletion of Pole3 and Pole4 or mutations at the HBD of Mcm2 grow normally and maintain stemness. As parental histones with their histone modifications are the blueprint for the recapitulation of the epigenetic landscape during cell division (Corpet and Almouzni, 2009), we investigated the roles of these parental histone chaperone proteins during differentiation of mouse ES cells. We found that Mcm2, relying on its histone binding domain, promotes mouse ESC differentiation. The Mcm2-2A mutant impairs ES cell differentiation accompanied by dramatic changes of histone modifications, gene expression and chromatin accessibility during differentiation. Furthermore, we found that Mcm2 localizes at transcription starting sites and that this localization is dramatically reduced in Mcm2-2A cells. Finally, the reduction of Mcm2 localization in Mcm2-2A mutant cells correlates with reduced chromatin accessibility at bivalent chromatin domains during differentiation. Together, these studies reveal a novel role for Mcm2 and its ability to bind H3-H4 in the differentiation of mouse ES cells.

## Results

### Mcm2, Pole3 and Pole4 are required for the differentiation of mouse embryonic stem cells

CAF-1, Asf1a and HIRA are histone chaperones involved in deposition of newly synthesized H3-H4, whereas Mcm2, Pole3 and Pole4 are involved in parental histone transfer and recycling (Serra-Cardona and Zhang, 2018). It is known that CAF-1, Asf1a and HIRA play essential roles during stem cell differentiation (Banaszynski *et al*., 2013; Cheloufi *et al*., 2015; Cheng *et al*., 2019; Gao *et al*., 2018). We therefore asked whether histone chaperones involved in parental histone transfer are also important in this process. To do this, we firstly monitored the formation of embryoid bodies (EBs), which mimics the formation of three germ layers *in vitro*, in Pole3 KO, Pole4 KO, Mcm2-2A single and double mutant mouse embryonic stem cells (Li *et al*., 2020) (Figure 1—figure supplement 1A). Briefly, three dimensional colonies (EBs) were formed in hanging drops without leukemia inhibiting factor (LIF) for 3 days. Then, EBs were cultured in suspension and collected at different times during differentiation for the evaluation of morphology and expression of selected genes involved in stemness and lineage specification (Figure 1A).

**Figure 1:**
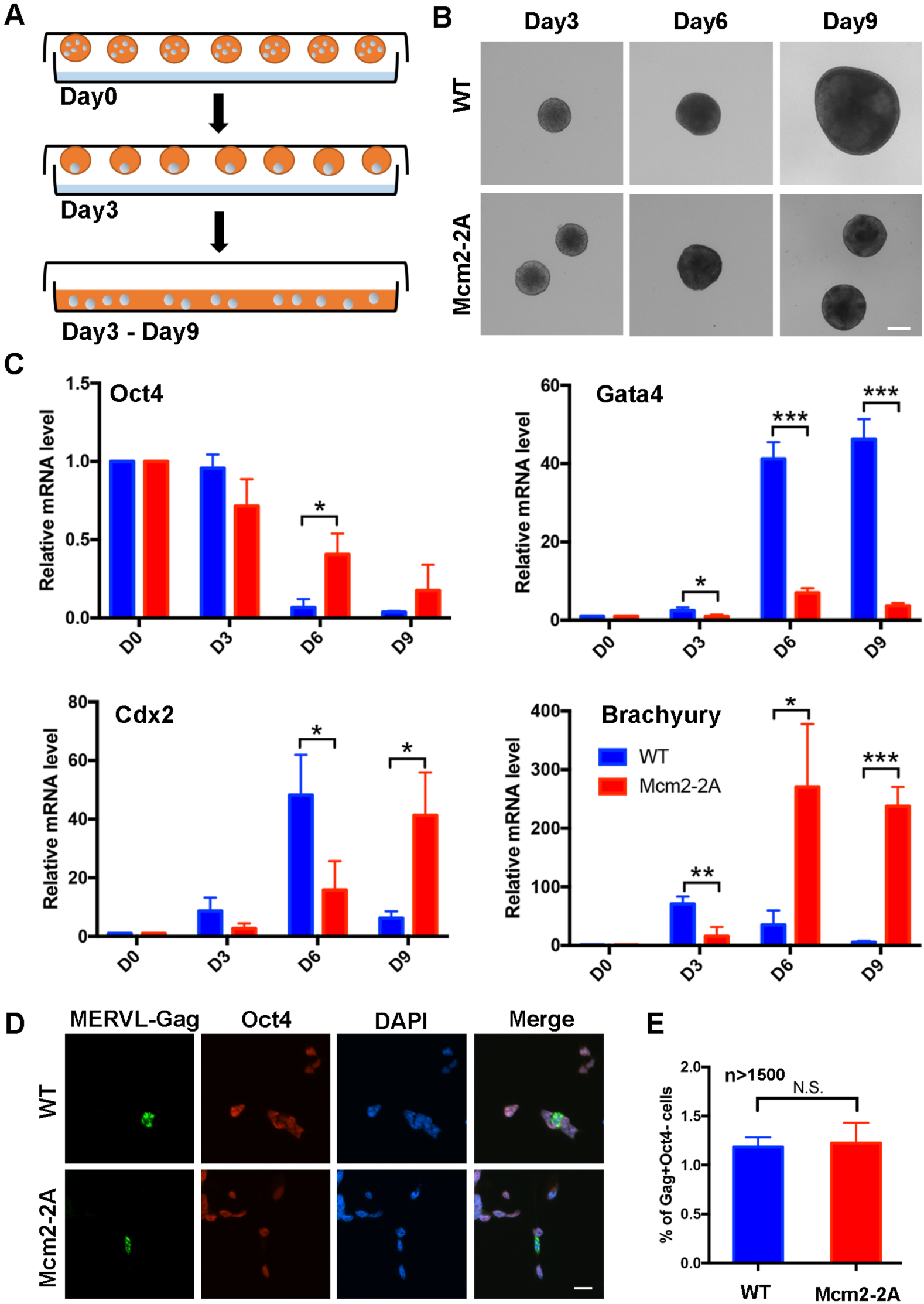
Mcm2-2A mutation in mouse ES cells impairs differentiation. See also Figure 1-source data 1. (A) A diagram showing the embryoid body (EB) formation assay. (B) Representative images of WT and Mcm2-2A cells during the process of EB formation. Scale bar: 20 µm. (C) RT-PCR analysis of the expression of *Oct4* (a gene involved in pluripotency) and three lineage-specific genes in WT and Mcm2-2A cells during EB formation. GAPDH was used for normalization. Data are presented as means ± SD from three independent experiments. (D) Representative immunofluorescence images for detection of Oct4 and MERVL-Gag proteins in WT and Mcm2-2A mouse ESCs. Scale bar: 20 µm. (E) Quantification of Gag+ Oct4-cells in (D). At least 1500 cells were counted for each cell line. Data are presented as means ± SD from three independent experiments. Statistical analysis in C and E was performed by two-tailed unpaired Student *t* test (\**P* < 0.05; \*\**P* < 0.01; \*\*\**P* < 0.001, N.S., no significant difference).

We observed that all mutants exhibited reduced EB size at day 9 compared to wild type cells, with Mcm2-2A, Mcm2-2A Pole3 KO, Mcm2-2A Pole4 KO double mutant cells exhibiting dramatic defects based on morphology (Figure 1B, Figure 1—figure supplement 1A). Similarly, we observed that the silencing of *Oct4,* a gene involved in pluripotency, was compromised in all the mutant cells, with the Mcm2-2A Pole3 KO and Mcm2-2A Pole4 KO double mutant cells showing larger effects than either single mutant alone (Figure 1C, Figure 1—figure supplement 1B). Finally, the transcription of several lineage specific genes, representing three germ layers, was delayed in all the mutant cells, with the double mutants showing the strongest defects (Figure 1C, Figure 1—figure supplement 1B). Together, these studies indicate that Mcm2-2A, Pole3 KO and Pole4 KO mutations, while having little impacts on the stemness of mouse ES cells, have a profound effect on their differentiation.

Depletion of CAF-1 results in an increase in totipotent 2-cell-like cells, which are marked by reduced expression of Oct4 and increased expression of the endogenous retrovirus MERVL (Ishiuchi *et al*., 2015). We thus assessed whether Mcm2-2A mutants altered cellular plasticity in ES cells by analyzing the expression of Oct4 and MERVL-Gag using immunofluorescence. We found that Mcm2-2A mutant and wild type cells showed similarly low frequencies of 2C-like cells (Figure 1D, E), indicating that ES cell plasticity is not altered in Mcm2-2A mutant cells. Together, these results suggest that the ability of Mcm2 to bind histone H3-H4 plays an important role during mouse ES cell differentiation.

### Mcm2-2A mutation compromises mouse ES cell differentiation into neural lineages

To further study the function of Mcm2 and Pole4 during ES cell differentiation, we differentiated wild type, Mcm2-2A and Pole4 KO single and double mutant ES cells into neural lineage cells as previously described (Gao *et al*., 2018). We then compared expression levels of two pluripotency genes (*Oct4* and *Nanog*) and two neural lineage specific genes (*Sox21* and *Pax6*) in WT, Mcm2-2A, Pole4 KO, and Mcm2-2A Pole4 KO double mutant cells. Consistent with the EB formation assays, Mcm2-2A and double mutant (Mcm2-2A and Pole4 KO) cells exhibited compromised silencing of Oct4 and Nanog and showed defects in the induction of Sox21 and Pax6 expression (Figure 2A, Figure 2—figure supplement 1A). In contrast, Pole4 KO had no significant effects on the expression of pluripotency genes or neural lineage genes compared with WT cells during neural differentiation (Figure 2—figure supplement 1A).

**Figure 2:**
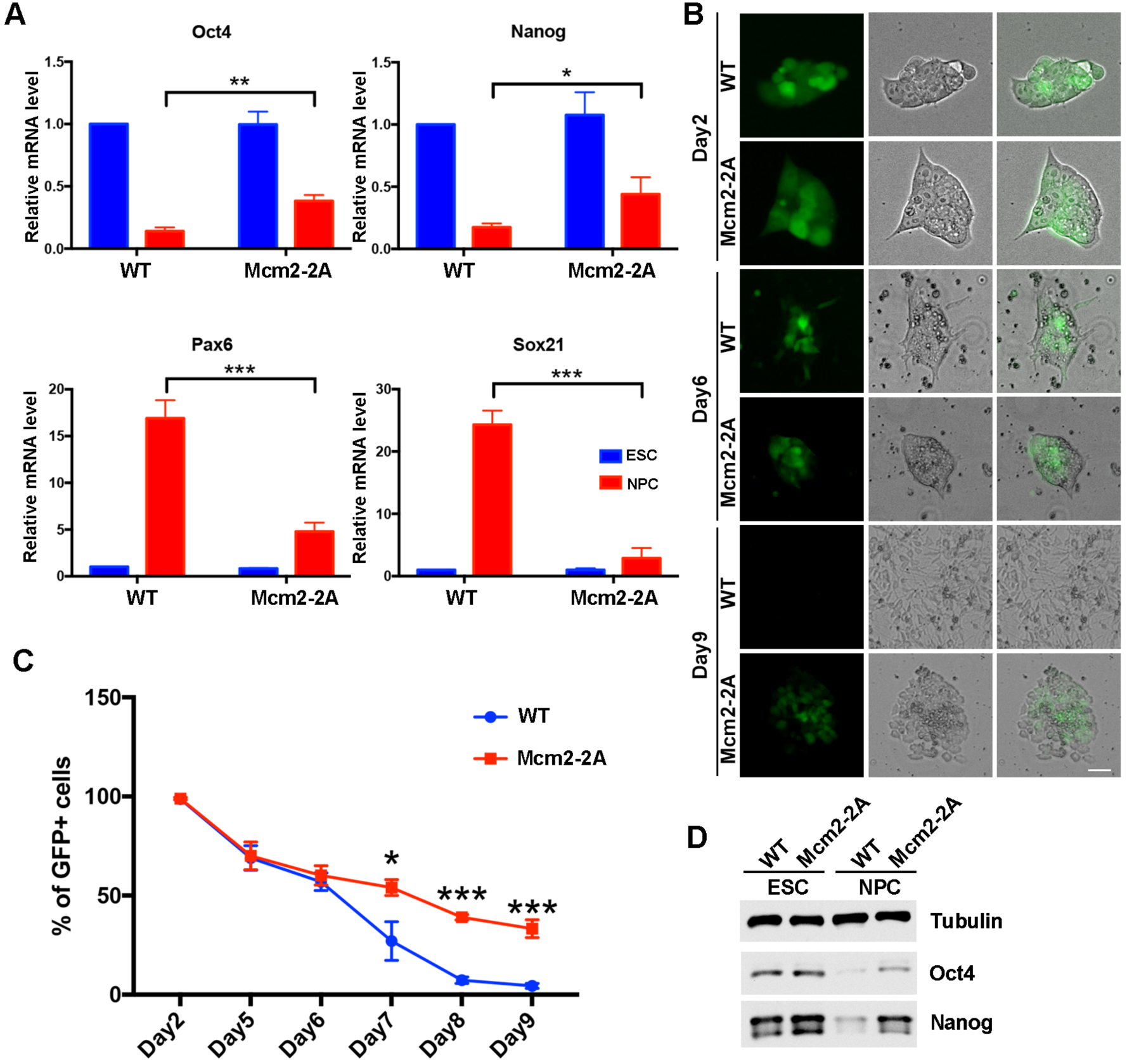
Mcm2 is required for neural differentiation of mouse ES cells. See also Figure 2-source data 1 and Figure 2-source data 2. (A) RT-PCR analysis of expression of two genes involved in pluripotency (*Oct4*, *Nanog*) and two neural lineage specific genes (*Pax6*, *Sox21*) in WT and Mcm2-2A cells during neural differentiation. GAPDH was used for normalization. Data are presented as means ± SD from three independent experiments. (B) Representative images of WT and Mcm2-2A cells during neural differentiation. The expression of EGFP is driven by the Oct4 distal enhancer. Scale bar: 20 µm. (C) FACS analysis of the percentage of GFP+ cells in WT and Mcm2-2A cells during neural differentiation. Data are presented as means ± SD from three independent experiments. Statistical analysis in A and C was performed by two-tailed unpaired Student *t* test (\**P* < 0.05; \*\**P* < 0.01; \*\*\**P* < 0.001). (D) WB analysis of Oct4 and Nanog in WT and Mcm2-2A cells of both ESCs and NPCs. Tubulin was used as a loading control.

Because Mcm2-2A mutant ES cells showed consistent defects in differentiation in both EB formation and neural differentiation assays, we focused the remaining studies on this mutant. First, we monitored morphology changes and silencing of the EGFP reporter gene driven by the Oct4 distal enhancer during neural differentiation in WT and Mcm2-2A cells (Hotta et al., 2009). The silencing of EGFP reporter and the cell morphologies were comparable between WT and Mcm2-2A cells from day 2 to day 6 (Figure 2B, C; Figure 2—figure supplement 1B). Interestingly, from day 7 to day 9, we observed a dramatic loss of GFP expression in WT cells, whereas ∼30% of Mcm2-2A cells preserved GFP signals at day 9 (Figure 2B, C; Figure 2—figure supplement 1B). Accordingly, we detected higher expression level of Oct4 and Nanog proteins in Mcm2-2A cells compared to wild type cells after differentiation (Figure 2D). Finally, Mcm2-2A cells upon differentiation did not exhibit neural cell morphology like WT cells (Figure 2B), providing additional support to the idea that the Mcm2-2A mutation compromises the differentiation of mouse ES cells.

### Mcm2-2A mutation perturbs transcription globally during differentiation

To understand how the Mcm2-2A mutation affects ES cell differentiation, we used RNA sequencing (RNA-seq) and analyzed the transcriptome of WT and Mcm2-2A mouse ES cells (ESCs) as well as neural precursor cells (NPCs) collected on day 9 of neural differentiation. In general, the Mcm2-2A mutation caused greater gene expression changes in NPCs than in ES cells, with 54 differentially expressed genes (DEGs) between wild type and Mcm2-2A ES cells and 609 DEGs between wild type and Mcm2-2A NPCs (false discovery rate (FDR)<0.01, |LFC|>1) (Figure 3A). Therefore, the Mcm2-2A mutation did not affect gene transcription in ES cells dramatically, providing an explanation for the observation that the Mcm2-2A mutation does not affect stemness or proliferation of mouse ES cells dramatically. Inspection of RNA-seq data confirmed that the silencing of pluripotency gene *Oct4* as well as the induction of two neural lineage genes (*Sox21* and *Pax6*) in Mcm2-2A NPCs was compromised compared to wild type NPCs (Figure 3B). Unsupervised cluster analysis separated the 648 DEG genes identified in Mcm2-2A cells into 4 groups (Figure 3C). Gene ontology (GO) analysis of 352 down-regulated genes in Mcm2-2A NPCs were enriched in neurodevelopment, such as forebrain/hindbrain development and neural nucleus development (Figure 3D), whereas 238 up-regulated genes were related to stem cell population maintenance (Figure 3E). Thus, the Mcm2-2A mutation induces aberrant expression of genes, including both up-regulated and down-regulate genes, during neural differentiation, which in turn contributes to the differentiation defects observed in the mutant cells.

**Figure 3:**
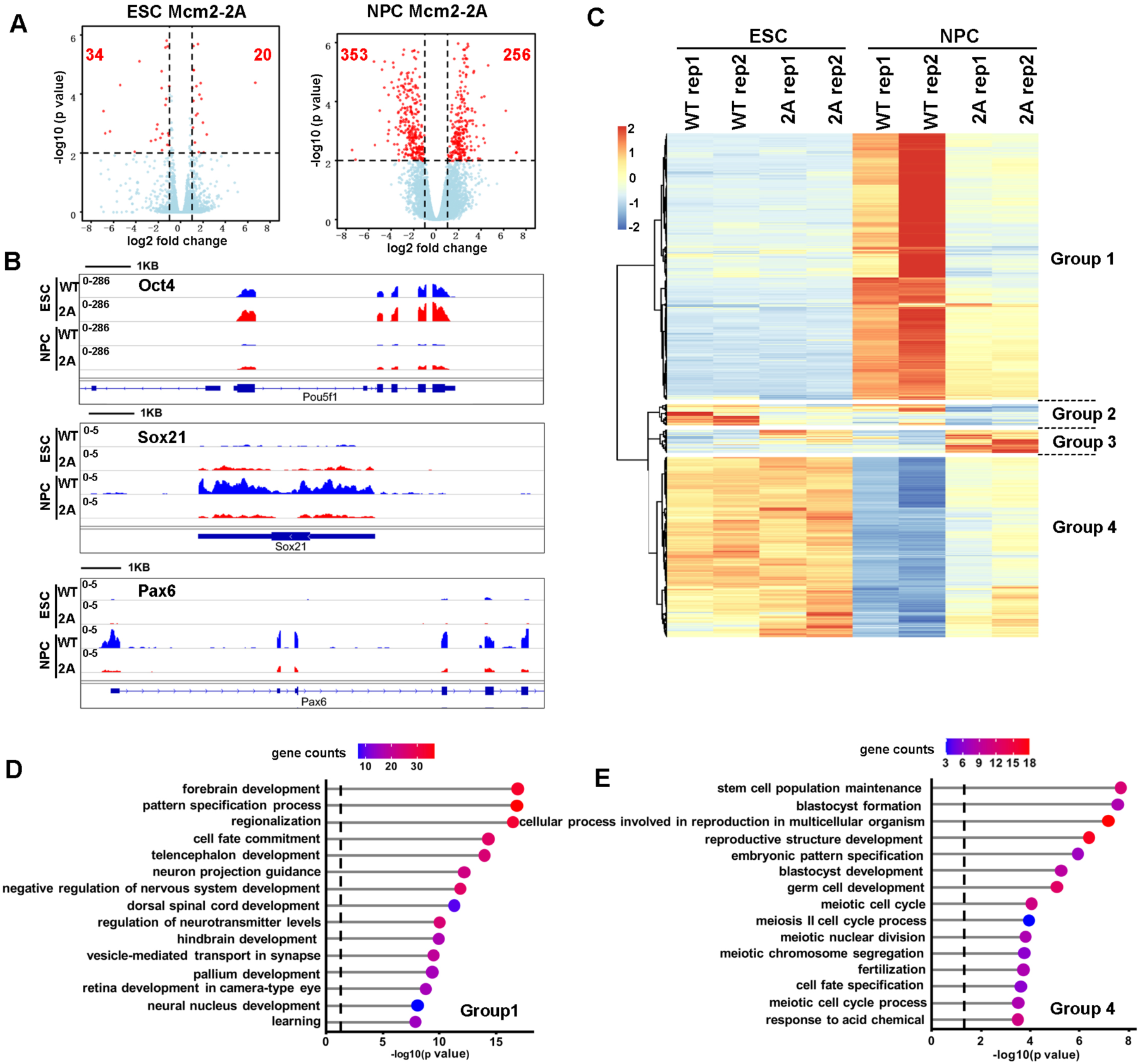
Effects of Mcm2-2A mutation on gene transcription in ES cells and in NPCs. (A) Volcano plot of differentially expressed genes (DEGs) between wild type and Mcm2-2A mutant ES cells (left) and neural precursor cells (NPCs, right) from two independent replicates, with numbers of significantly up-regulated and down-regulated genes (red dots, P<0.01, |log2 fold change|>1) shown. (B) RNA-seq tracks showing the sequence read density at Oct4, Sox21 and Pax6 locus in WT and Mcm2-2A ES cells and NPCs. (C) The hierarchical clustering analysis of the differentially expressed genes between WT and Mcm2-2A cells. (D and E) GO analysis of the group 1 and group 4 genes in (C) with the top 15 significant GO terms and *P* value displayed.

### Mcm2-2A mutation disrupts the epigenetic landscape in differentiated cells

During differentiation, histone modification landscapes are rewired (Mikkelsen *et al*., 2007). For instance, bivalent domains that are enriched with both active markers (H3K4me3) and silencing markers (H3K27me3), which are associated with lineage specific genes, are resolved either through the removal of repressive marker H3K27me3 for gene activation or removal of active marker H3K4me3 for gene silencing (Voigt et al., 2013). In addition, pluripotency genes are silenced through gain of H3K27me3 and loss of H3K4me3 at promoters (Bernstein *et al*., 2006; Harikumar and Meshorer, 2015). Therefore, we analyzed the impact of the Mcm2-2A mutation on the total levels of H3K27me3 and H3K4me3 during neural differentiation. We observed that in both wild type and Mcm2-2A mutant NPCs, the H3K27me3 level was reduced compared to their corresponding ES cells, with a modest reduction of H3K27me3 in Mcm2-2A NPCs compared to WT NPCs (Figure 4A). In contrast, the Mcm2-2A mutation did not have an effect on H3K4me3 levels during differentiation (Figure 4A). Next, we determined genome-wide distributions of H3K4me3 and H3K27me3 in wild type and Mcm2-2A ES cells as well as NPCs using H3K4me3 and H3K27me3 CUT&RUN (Figure 4B). Similar to the transcriptome changes, the effects of Mcm2-2A on the chromatin binding of H3K4me3 or H3K27me3 in ES cells were much less pronounced than that in NPCs (Figure 4B). Globally, H3K4me3 CUT&RUN density was similar in Mcm2-2A ESCs compared to wild type ESCs at TSS (transcription starting site) of both down-regulated and up-regulated genes in Mcm2-2A ESCs (Figure 4—figure supplement 1A, B).

**Figure 4:**
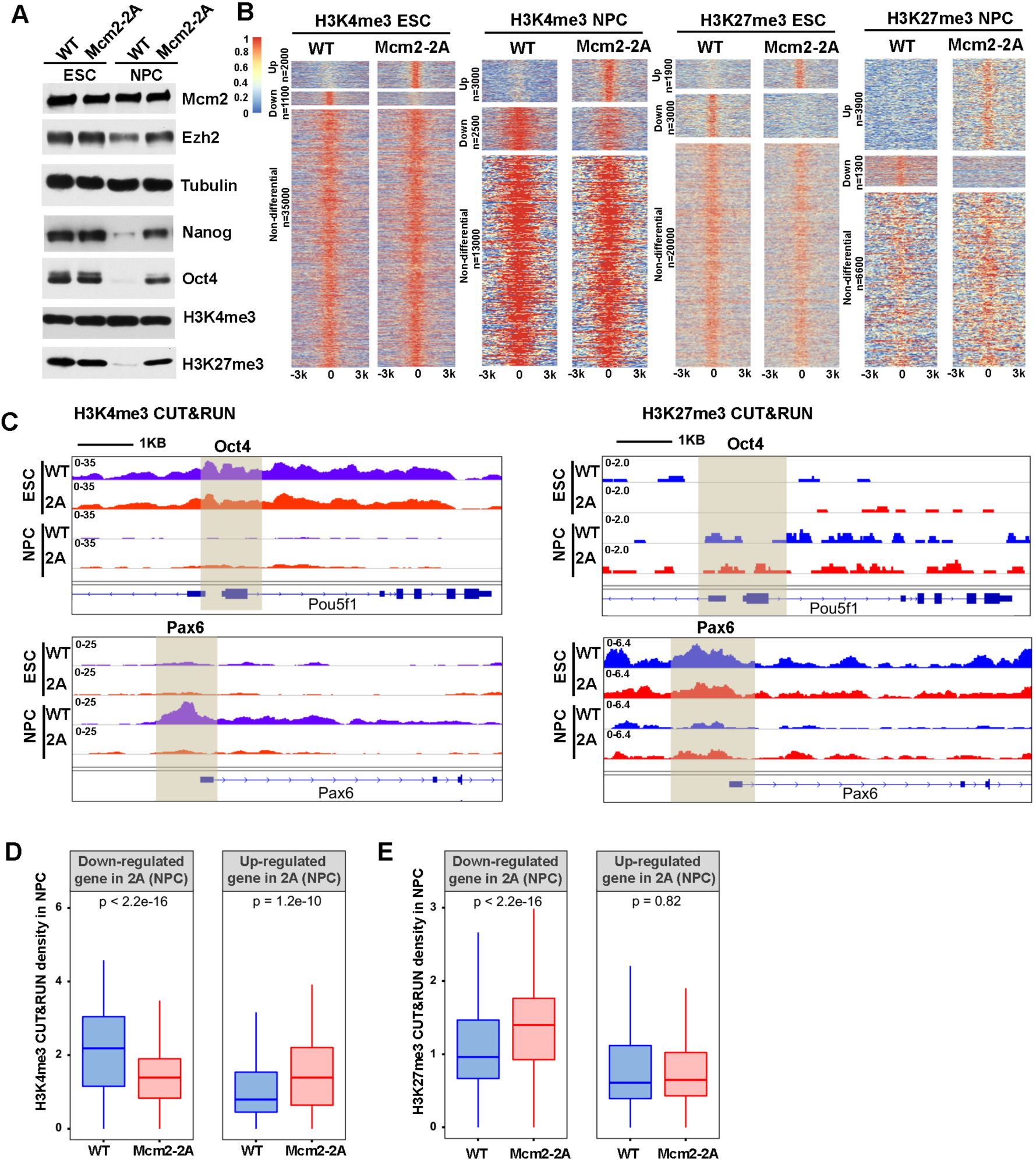
Mcm2 mutant affects dynamic changes in H3K4me3 and H3K27me3 during differentiation. See also Figure 4-source data 1. (A) WB analysis of Mcm2, Ezh2, Nanog, Oct4, H3K4me3 and H3K27me3 expression in WT and Mcm2-2A cells of both ES cells and NPCs. Tubulin was used as loading control. (B) Heatmaps surrounding CUT&RUN peaks of H3K4me3 and H3K27me3 [-3k, 3k] in WT and Mcm2-2A ES cells and NPCs. n: peak number. (C) H3K4me3 and H3K27me3 CUT&RUN sequencing density surrounding Oct4 and Pax6 loci in WT and Mcm2-2A ES cells and NPCs. Shadows indicate CUT&RUN peak signals around the TSS. (D and E) Average of H3K4me3 (D) and H3K27me3 (E) CUT&RUN density in WT and Mcm2-2A NPCs at the promoters ([-3k, 3k] of TSS) of down-regulated and up-regulated genes in Mcm2-2A NPCs (Figure 3A, right). The Y-axis represents the log2 ratio of CUT&RUN density (RPKM). The *P* values were calculated using Wilcoxon signed-rank test. The average of two independent replicates are shown.

In NPCs, we observed that H3K4me3 CUT&RUN density at the *Oct4* gene promoter was dramatically reduced compared to wild type ES cells, whereas the level of H3K4me3 at the promoter of *Pax6* was dramatically increased in WT NPCs compared to WT ESCs. These results are consistent with the silencing of Oct4 and induction of Pax6 during differentiation in wild type ESCs. In Mcm2-2A mutant cells, both the reduction of H3K4me3 at the promoter of *Oct4* and the increase of this mark at the promoter of *Pax6* were compromised, consistent with the compromised silencing of Oct4 and the induction of Pax6 in Mcm2-2A mutant cells during differentiation (Figure 4C). Furthermore, the average levels of H3K4me3 at the TSS of down-regulated genes in Mcm2-2A mutant NPCs were significantly lower than those of WT NPCs and higher at the TSS of up-regulated genes (Figure 4D, Figure 4—figure supplement 1C). Together, these results indicate that the gene expression changes in Mcm2-2A mutant NPCs correlate with changes in H3K4me3 levels at gene promoters.

While we observed far more increased H3K27me3 peaks than reduced H3K27me3 peaks in Mcm2-2A mutant NPCs compared to wild type NPCs (Figure 4B), the changes in gene expression in Mcm2-2A NPCs also correlated with changes H3K27me3. Specifically, H3K27me3 density at the *Pax6* promoter was higher in Mcm2-2A than wild type NPCs, whereas H3K27me3 density at the *Oct4* promoter were similar (Figure 4C). On average, H3K27me3 density at the TSS of down-regulated genes in Mcm2-2A mutant NPCs was significantly higher than WT NPCs. In contrast, H3K27me3 density at the TSS of up-regulated genes was similar between wild type and Mcm2-2A NPCs (Figure 4E, Figure 4—figure supplement 1D), suggesting that the reduced expression of genes in Mcm2-2A NPCs likely was due to the retention of this repressive mark during differentiation. Collectively, these results suggest that the gene expression alterations in Mcm2-2A mutant NPCs compared to wild type NPCs correlate with the changes of both H3K4me3 and H3K27me3 levels at gene promoters.

### Mcm2 is enriched at actively transcribed regions in both ES cells and in NPCs

To further explore the mechanisms underlying the transcriptome and epigenetic changes in Mcm2-2A mutant cells, we performed Mcm2 CUT&RUN in both wild type and Mcm2-2A ES cells and NPCs. Initial studies of Mcm2 CUT&RUN using antibodies against Mcm2 or against the Flag epitope fused to Mcm2 and Mcm2-2A proteins in ES cells (Xu et al., 2022) indicate that Mcm2 antibody and Flag antibody CUT&RUN profiles, each of which had two independent repeats, were highly correlated with each other (Figure 5—figure supplement 1A, B), demonstrating the reproducibility and reliability of Mcm2 CUT&RUN datasets. Using *P*=0.0001 as a cutoff, we identified 13742 Mcm2 CUT&RUN peaks in wild type ES cells. We then aligned and clustered these Mcm2 CUT&RUN peaks based on their overlaps with H3K4me3 and H3K27me3 CUT&RUN signals and ATAC-seq peaks in ES cells (Figure 5A). We observed that the majority of Mcm2 CUT&RUN peaks were enriched with H3K4me3 CUT&RUN signals and ATAC-seq peaks. A small fraction of Mcm2 CUT&RUN peaks were found at bivalent chromatin domains (H3K4me3+ and H3K27me3+), with far fewer peaks co-localizing with H3K27me3 markers (H3K4me3-, H3K27me3+) in ES cells (Figure 5A, left). We also performed similar analysis on 2686 of Mcm2 peaks identified in wild type NPCs and observed that almost all the Mcm2 peaks co-localized with H3K4me3 and ATAC-seq peaks (H3K4me3+, H3K27me3-) (Figure 5A, right). These results indicate that Mcm2 binding to chromatin, like histone modifications, is rewired during differentiation. Moreover, Mcm2 CUT&RUN signals were enriched at the transcription starting site (TSS) of highly transcribed genes compared to the TSS of lowly transcribed genes in both ES cells and NPCs (Figure 5B, Figure 5—figure supplement 1C), indicating that the majority of Mcm2 localizes at actively transcribed regions. Since Mcm2 is an important subunit of the Mcm2-7 complex (Tye, 1999), we asked if Mcm2 peaks are enriched at DNA replication origins in mouse ES cells (Li *et al*., 2020). Mcm2 CUT&RUN peaks exhibited low densities at origins (Figure 5—figure supplement 1D) and were highly enriched at ∼316 kb away from the center of origins (Figure 5—figure supplement 1E). It is known that DNA replication initiates stochastically in mammalian cells (Prioleau and MacAlpine, 2016), which may contribute to our inability to detect the enrichment of Mcm2 at replication origins. Nonetheless, these results indicate that Mcm2 may also function in gene transcription in a manner independent of its role in DNA replication.

**Figure 5:**
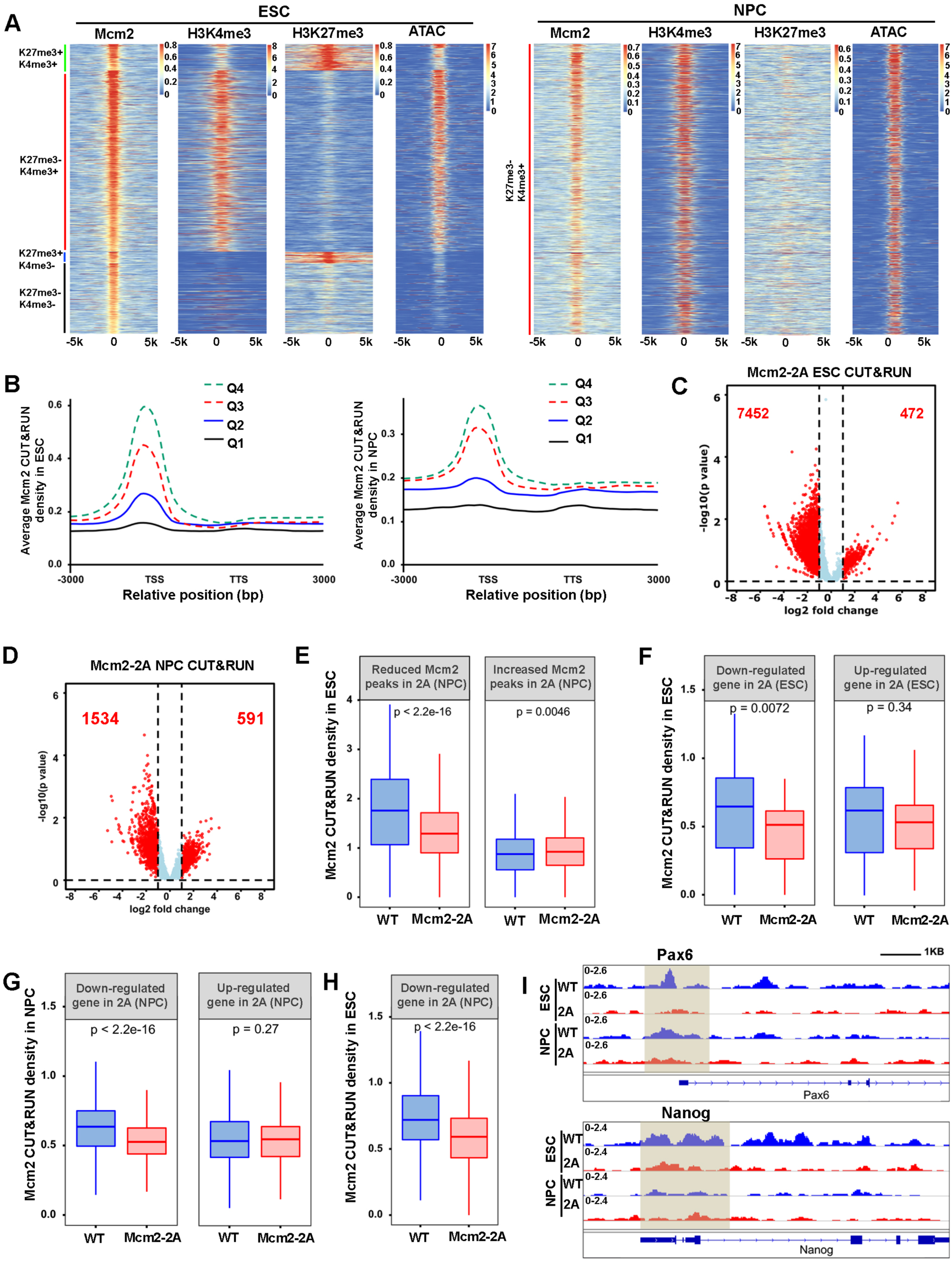
Mcm2 chromatin localization largely depends on its ability to bind H3-H4. (A) Representative heatmaps of Mcm2, H3K4me3, H3K27me3 CUT&RUN and ATAC-seq peaks in WT ES cells (left) and NPCs (right). The density of H3K4me3 and H3K27me3 CUT&RUN and ATAC-seq surrounding Mcm2 CUT&RUN peaks [-5k, 5k] was calculated. (B) Density profiles of Mcm2 CUT&RUN (RPM, reads per million) surrounding transcription starting sites (TSS) and transcription termination sites (TTS). Genes were separated into 4 groups based on their expression in mouse ES cells or NPCs (Q1 = lowest quartile, Q4 = highest quartile). (C and D) Volcano plot of differential Mcm2 protein CUT&RUN peaks between WT Mcm2 and Mcm2-2A ES cells (C) and NPCs (D) from two independent replicates, with the total number of significantly up-regulated and down-regulated peaks (|log2 fold change|>1) shown. (E) Mcm2 CUT&RUN density in WT and Mcm2-2A ES cells at the reduced and increased Mcm2 CUT&RUN peaks in Mcm2-2A NPCs identified in Figure 5D. (F) Mcm2 CUT&RUN density in WT and Mcm2-2A ES cells at the promoters ([-3k, 3k] of TSS) of down-regulated and up-regulated genes in Mcm2-2A mutant ES cells (ESCs) identified in Figure 3A, left. (G) Mcm2 CUT&RUN density in WT and Mcm2-2A NPCs at the promoters ([-3k, 3k] of TSS) of down-regulated and up-regulated genes in Mcm2-2A mutant NPCs (Figure 3A, right). (H) Mcm2 CUT&RUN density in WT and Mcm2-2A ES cells at the promoters ([-3k, 3k] of TSS) of down-regulated genes in Mcm2-2A mutant NPCs (Figure 3A, right). (E-H) The Y-axis represents the log2 ratio of CUT&RUN density (RPKM), with *P* values calculated using Wilcoxon signed-rank test from two independent replicates. (I) Snapshots displaying Mcm2 CUT&RUN density at Oct4 and Pax6 loci of WT and Mcm2-2A ES cells and NPCs. One representative result from two independent replicates is shown. Shadows indicate CUT&RUN signals around the TSS.

Next, we compared Mcm2 WT and Mcm2-2A mutant CUT&RUN profiles in both ES cells and NPCs. We observed far more reduced Mcm2 CUT&RUN peaks than increased peaks in both Mcm2-2A mutant ES cells and NPCs cells compared with their WT counterparts (Figure 5C, D; Figure 5—figure supplement 1B). Most of these Mcm2 reduced peaks occurred at active promoters and bivalent chromatin regions, whereas increased peaks showed no specific enrichment (Figure 5—figure supplement 1F). Because of dramatic changes of Mcm2 localization in Mcm2-2A ES cells and NPCs, we first asked whether the changes of Mcm2 CUT&RUN peaks in ES cells correlated with those in NPCs. We observed that at the reduced Mcm2 peaks in Mcm2-2A NPCs, Mcm2 CUT&RUN density in Mcm2-2A ES cells was also significantly lower than in wild type ES cells, whereas at increased Mcm2-2A CUT&RUN peaks in Mcm2-2A NPCs, Mcm2 density in Mcm2-2A ES cells was higher than in wild type ES cells (Figure 5E). Together, these results suggest that alterations in chromatin binding of Mcm2-2A mutant proteins at ES cells likely contribute to its altered chromatin binding in NPCs.

Next, we asked whether the changes in Mcm2 binding in ES cells and NPCs correlated with changes in gene expression in ES cells and in NPCs. We found that the average level of Mcm2-2A density at the TSS of down-regulated genes in ES cells was significantly lower than that in WT ES cells, but Mcm2 density at up-regulated genes was similar between wild type and Mcm2-2A mutant ES cells (Figure 5F). Similar results were obtained in NPCs (Figure 5G). In addition, Mcm2 CUT&RUN density in Mcm2-2A mutant ES cells was reduced at the down-regulated genes in NPCs (Figure 5H), as exemplified at the promoter of neural lineage gene *Pax6*, where Mcm2 CUT&RUN density was down-regulated in mutant NPCs (Figure 5I). This finding suggests that the reduced association of Mcm2 with chromatin in Mcm2-2A ES cells likely contributes to the reduced expression of these genes in NPCs. Taken together, these results indicate that Mcm2 localizes at the TSS of actively transcribed regions in both ES cells and NPCs and that the histone binding of Mcm2 is important for its chromatin binding in ES cells and NPCs. Furthermore, the reduced chromatin binding of Mcm2-2A in ES cells, while having minor effects on gene expression, may have marked effects on gene expression and Mcm2 binding during differentiation.

### Mcm2 facilitates chromatin accessibility during mouse ES cell differentiation

The landscape of chromatin accessibility dynamically changes during the development (Trevino et al., 2020). Given Mcm2’s role in recycling parental histones (Huang *et al*., 2015; Li *et al*., 2020; Petryk *et al*., 2018; Xu *et al*., 2022) and the high correlation of Mcm2 CUT&RUN peaks with ATAC-seq signals in ES cells (Figure 5A), we analyzed chromatin accessibility in WT and Mcm2-2A mutant ES cells and NPCs using ATAC-seq. The ATAC-seq repeats in both ES cells or NPCs were highly correlated with each other (Figure 6—figure supplement 1A, B). Volcano plot analysis and genome-wide correlation revealed that in ES cells, the Mcm2-2A mutation did not markedly change ATAC-seq profiles (Figure 6—figure supplement 1A, C), In contrast, ATAC-seq profiles in Mcm2-2A mutant NPCs were dramatically altered compared to wild type NPCs (Figure 6—figure supplement 1B, D). These results are consistent with the observations that the Mcm2-2A mutation had little impact on gene expression in ES cells, but markedly altered gene expression in NPCs. Importantly, ATAC-seq signals, which reflect chromatin accessibility, in NPCs were highly correlated with gene expression levels. For instance, we observed an increase and a reduction of ATAC-seq signals at the promoters of *Oct4* and *Pax6*, respectively, in Mcm2-2A NPCs (Figure 6A). Moreover, the average level of ATAC-seq signals in Mcm2-2A mutant NPCs was significantly lower than WT NPCs at the TSS of down-regulated genes in NPCs and higher than WT NPCs at up-regulated genes (Figure 6B, Figure 6—figure supplement 2A). These results are consistent with the idea that chromatin accessibility, as detected by ATAC-seq, is linked to gene transcription.

**Figure 6:**
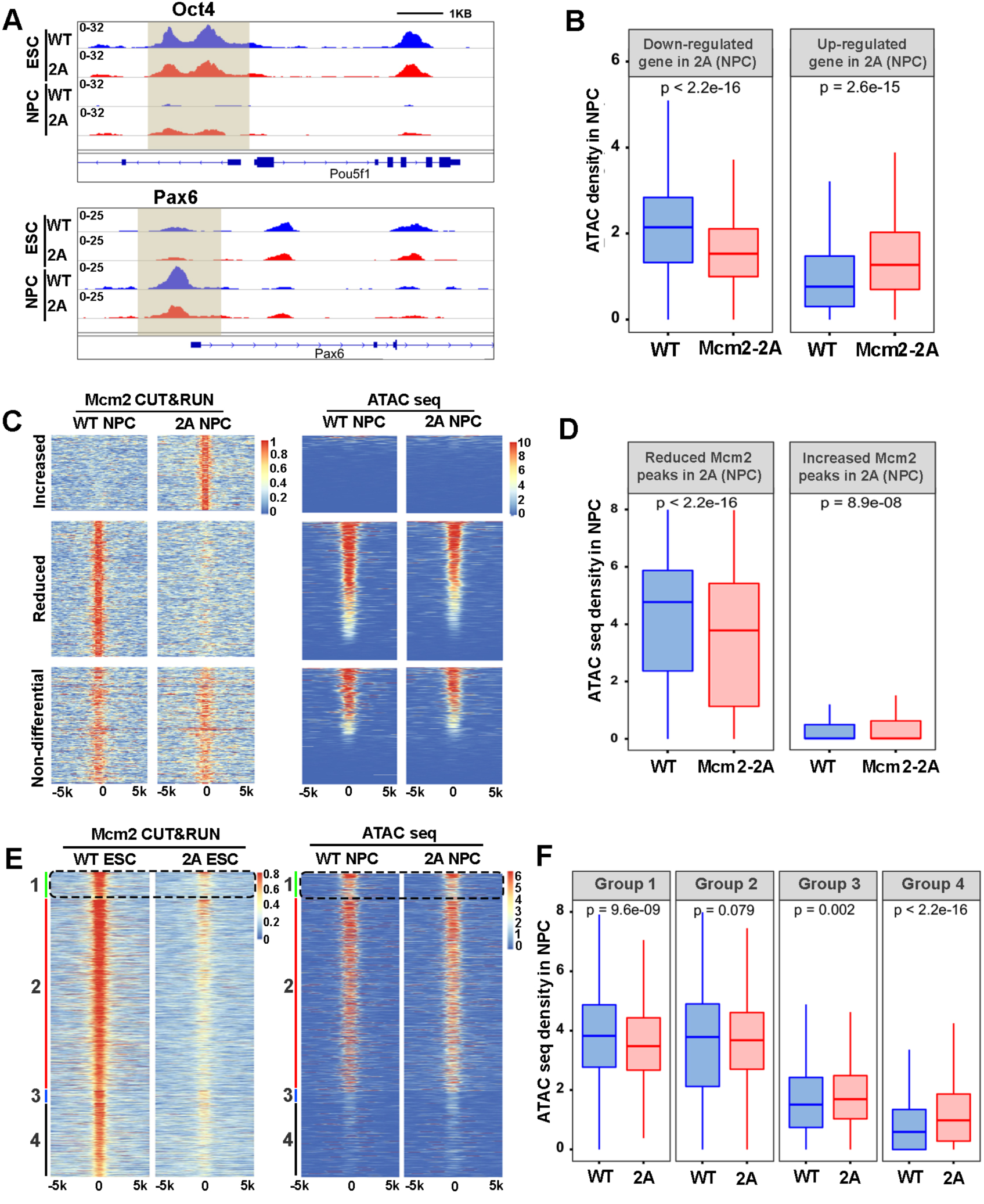
Mcm2 facilitates chromatin accessibility during mouse ES cell differentiation. (A) ATAC-seq tracks displaying ATAC-seq density at the Oct4 and Pax6 loci in WT and Mcm2-2A ES cells and NPCs. Shadows indicate CUT&RUN signals around the TSS. (B) ATAC-seq density in WT and Mcm2-2A NPCs at the promoters ([-3k, 3k] of TSS) of down-regulated and up-regulated genes in Mcm2-2A NPCs (Figure 3A, right). (C) Heatmap of ATAC-seq density (left) surrounding Mcm2 CUT&RUN peaks (right, |log2 fold change|>1) in WT and Mcm2-2A NPCs. One representative result from two independent replicates is shown. (D) The average of ATAC-seq density in wild type and Mcm2-2A NPCs at the reduced and increased Mcm2 CUT&RUN peaks in Mcm2-2A NPCs shown in C. (E) Representative heatmap of ATAC-seq peaks in wild type and Mcm2-2A NPCs at four groups of Mcm2 peaks identified in wild type ES cells (Figure 5A) based on their co-localization with H3K27me3 and H3K4me3: Group 1: H3K4me3+ and H3K27me3+ (bivalent domain); Group 2: H3K4me3+ and H3K27me3-(active promoters); Group 3: H3K4me3- and H3K27me3+ (repressive promoters); and Group 4: H3K4me3- and H3K27me3-. (F) Average ATAC-seq density in WT and Mcm2-2A NPCs at each of the four groups shown in E. (B, D and F) The Y-axis represents the log2 ratio of ATAC seq density (RPKM). The *P* values were calculated using Wilcoxon signed-rank test. The average of two independent replicates is shown.

Next, we explored the correlation between Mcm2 binding and ATAC-seq density. Based on Figure 5C and 5D, we first separated Mcm2-2A CUT&RUN peaks in ES cells and in NPCs into three categories (increased, reduced and non-differential Mcm2 peaks compared to wild type Mcm2 ES cells and NPCs) and calculated ATAC-seq density at each group of Mcm2 peaks. We observed a dramatic reduction of ATAC-seq density in Mcm2-2A mutant NPCs within the reduced Mcm2 peak group in NPCs, along with a slight increase of ATAC-seq density in Mcm2-2A mutant NPCs at increased Mcm2 peak groups (Figure 6C, D). Of note, ATAC-seq signals at increased Mcm2 peaks were very low (Figure 6C, D). In ES cells, ATAC-seq density in Mcm2-2A mutant cells was also reduced at down-regulated Mcm2 peaks, but to a far lesser extent than in NPCs (Figure 6—figure supplement 2B, C), implying a dominant effect of Mcm2 distribution on chromatin accessibility at differentiated cells.

Since Mcm2 binding in ES cells likely affects gene expression in NPCs (Figure 5), we further analyzed the correlation of Mcm2 binding in ES cells with chromatin accessibility. Because Mcm2 peaks in ES cells can be classified into four groups based on their co-localization with H3K27me3 and H3K4me3, we analyzed ATAC-seq signals at each group in wild type and Mcm2-2A ES cells and NPCs. In ES cells, while Mcm2 CUT&RUN density in Mcm2-2A mutant cells was reduced in all four groups (Figure 6—figure supplement 2D), ATAC-seq signals were slightly reduced in Group 2 (H3K4me3+, H3K27me3-) (Figure 6—figure supplement 2E, F). In NPCs, Mcm2 CUT&RUN density in Mcm2-2A mutant cells was reduced in both Group 1 and Group 2, with Group 1 showing a larger reduction than Group 2 (Figure 6—figure supplement 2G, H). Importantly, ATAC-seq signals in Mcm2-2A NPCs in Group 1 and Group 2 were significantly reduced compared to wild type NPCs, with a larger reduction in Group 1 than in Group 2, whereas ATAC-seq signals in Mcm2-2A NPCs were increased slightly in Group 3 and Group 4 (Figure 6E, F). As Mcm2 Group 1 peaks co-localize with bivalent chromatin domains (H3K4me3+ and H3K27me3+), these results indicate that a reduction of Mcm2 binding in Mcm2-2A mutant ES cells perturbs chromatin changes at bivalent domains during differentiation, and that Mcm2 binding at bivalent chromatin domains in ES cells is important for the resolution of these regions for subsequent gene activation during differentiation.

## Discussion

We and others have previously shown that Mcm2, Pole3 and Pole4, three replisome components first known for their roles in DNA replication, function in the transfer of parental H3-H4 following DNA replication in yeast and mouse ES cells (Gan *et al*., 2018; Li *et al*., 2020; Petryk *et al*., 2018; Xu *et al*., 2022; Yu *et al*., 2018). Remarkably, mouse ES cells with deletion of Pole3, Pole4 or with mutations at the histone binding motif of Mcm2 (Mcm2-2A) largely grow normally. Here, we found that these mutant cells all exhibit defects in differentiation, revealing a novel role of these proteins in ES differentiation.

Mcm2 is a subunit of MCM helicase consisting of Mcm2-7. The MCM helicase is loaded on chromatin at the G1/S transition and serves as the core of the CMG replicative helicase to unwind double stranded DNA for DNA synthesis during the S phase of the cell cycle (Tye, 1999). The N-terminus of Mcm2 contains a conserved histone binding motif that interacts with H3-H4 (Huang *et al*., 2015). Mutations at this histone binding motif (Mcm2-2A) that impair Mcm2’s ability to bind H3-H4 lead to a dramatic enrichment of parental H3-H4 at leading strands compared to lagging strands of DNA replication forks during early S phase of the cell cycle (Gan *et al*., 2018; Li *et al*., 2020; Petryk *et al*., 2018). Using two different *in vitro* differentiation assays, we observed that Mcm2-2A mutant mouse ES cells, while growing normally, showed dramatic defects during differentiation. Furthermore, we found that the Mcm2-2A mutation induces global changes in gene expression, chromatin accessibility and histone modifications in neural precursor cells compared to wild type NPCs. Together, these studies reveal a novel role of Mcm2, through its interaction with H3-H4, during differentiation.

Previously, we have shown that deletion of two histone chaperones involved in deposition of newly synthesized H3-H4, Asf1a and the p150 subunit of the CAF-1 complex, also impairs differentiation of mouse ES cells (Cheng *et al*., 2019; Gao *et al*., 2018). Compared to CAF-1 p150 KO and Asf1a KO cells, the Mcm2-2A mutation exhibited distinct defects in differentiation. p150 KO mutant ES cells show defects in both the silencing of pluripotent genes and induction of lineage specific genes during differentiation, with defects in the former much more pronounced (Cheng *et al*., 2019). In contrast, Asf1a KO ES cells minimally impact the silencing of pluripotent genes and dramatically affect the induction of lineage specific genes (Gao *et al*., 2018). Silencing of pluripotent genes and induction of lineage specific genes are both defective in Mcm2-2A cells, with the defects in the latter more pronounced. These results suggest that CAF-1, Asf1a and Mcm2 function in distinct aspects of chromatin regulation during differentiation.

During differentiation, the formation of heterochromatin at pluripotent genes and resolution of bivalent chromatin domains at lineage-specific genes are two obvious changes in chromatin (Bernstein *et al*., 2006; Mikkelsen *et al*., 2007). CAF-1 likely facilitates the formation of heterochromatin at pluripotent genes, whereas Asf1a promotes nucleosome disassembly at bivalent chromatin domains for the induction of lineage specific genes (Cheng *et al*., 2019; Gao *et al*., 2018). How, then, is Mcm2 involved in differentiation? We suggest four non-mutually exclusive models for Mcm2 to regulate chromatin dynamics and gene expression during differentiation. First, it is known that in Mcm2-2A mutant cells, the ability of MCM to bind Asf1 is also compromised (Huang *et al*., 2015). Therefore, it is possible that Mcm2 impacts chromatin through a mechanism similar to that of Asf1a. Supporting this idea, the induction of lineage of specific genes are defective in both Mcm2-2A and Asf1a KO mutant ES cells during differentiation. However, as mentioned above, Asf1a KO and Mcm2-2A ES cells show distinct defects during differentiation. Second, it is possible that Mcm2 is involved in the recycling of parental H3 during differentiation, which in turn contributes to dynamic changes in chromatin during differentiation. Recently, we have shown that Mcm2, Pole3 and Pole4 are involved in recycling both parental H3.3 and H3.1 in mouse ES cells following DNA replication (Xu *et al*., 2022). Furthermore, the N-terminus of Spt5 contains a conserved histone binding motif, and this binding is important for the recycling of parental H3 following gene transcription in budding yeast (Evrin et al., 2022). Currently, the fate of parental histones at different chromatin domains during mouse ES cell differentiation is largely unknown. It is possible that the ability of Mcm2 to recycle both parental H3.1 and H3.3 contributes to dynamic changes in chromatin states during differentiation. Third, it is possible that Mcm2, via its ability to bind H3-H4, regulates gene expression directly during differentiation. Supporting this idea, we have shown that Mcm2 localizes at gene promoters of actively transcribed genes in both ESCs and NPCs. Moreover, the reduced Mcm2 binding at bivalent chromatin domains in ES cells correlates with decreased chromatin accessibility at these regions in NPCs, suggesting that the ability of Mcm2 to bind H3-H4 is important to resolve bivalent chromatin domains during differentiation. Finally, MCM2-7 proteins are associated with RNA Pol II in *Xenopus* and HeLa cells (Snyder *et al*., 2009; Yankulov *et al*., 1999), and Mcm2 and Mcm5 are required for Pol II-mediated transcription (Snyder *et al*., 2009). Future studies are needed to address whether Mcm2’s role in gene transcription is linked to its ability to bind H3-H4.

We observed that the association of Mcm2-2A mutant proteins with chromatin is dramatically reduced in mouse ES cells compared to wild type ES cells. However, Mcm2-2A mutation did not affect chromatin accessibility and gene expression to a dramatic degree in mouse ES cells. We suggest two possible explanations for the minor phenotypes exhibited by Mcm2-2A mutant ES cells despite the dramatic reduction in chromatin association and defects in parental histone transfer during early S phase. First, it is known that excessive amounts of MCM2-7 helicase, greater than the number of replication origins, are loaded on chromatin during G1 phase of the cell cycle (Edwards et al., 2002; Lei et al., 1996). It is proposed that the excessive amount of MCM2-7 helicase, while not needed during unperturbed S phases of the cell cycle, helps fire dormant origins under replication stress (Woodward et al., 2006). We detected 13742 and 2686 Mcm2 CUT&RUN peaks, respectively, in wild type ES cells and NPCs, compared to 3339 and 1519 Mcm2 CUT&RUN peaks in Mcm2-2A ES and NPCs. From these numbers, it appears that excessive Mcm2 is found in mouse ES cells, with more Mcm2-2A mutant protein bound to chromatin in ES cells than wild type Mcm2 in NPCs. Therefore, we suggest that an excessive amount of Mcm2, and possibly Mcm2-7 complex in ES cells, safeguards not only DNA replication but also chromatin accessibility and gene expression in mouse ES cells. Second, we have shown recently in budding yeast that the asymmetric distribution of parental histones at replicating DNA strands in early S phase in Mcm2 mutant cells defective in parental histone transfer reduces dramatically during the cell cycle progression before cell division (Serra-Cardona et al., 2022). Therefore, it is likely that Mcm2-2A mutant ES cells restore the symmetric distribution of parental histones before cell division, minimizing the potential effects on mouse ES cells. Nonetheless, our results indicate that the Mcm2-2A mutation defective in histone binding impairs the differentiation of mouse ES cells, revealing a novel role of the ability of Mcm2 to bind H3-H4 during development.

## MATERIALS AND METHODS

### Materials availability statement

Further information and requests for resources and reagents should be directed to and will be fulfilled by the Lead Contact, Zhiguo Zhang.

### Cell culture and cell lines

The mouse E14 ES cell line (RRID:CVCL_C320) was kindly provided by Dr. Tom Fazzio (University of Massachusetts Medical School) and tested negative for mycoplasma. Cells were grown in DMEM (Corning) medium supplemented with 15% (v/v) fetal bovine serum (GeminiBio), 1% penicillin/streptomycin (Gibco), 1 mM sodium pyruvate (Gibco), 2 mM L-glutamine (Gibco), 1% MEM non-essential amino acids (Gibco), 55 µM β-Mercaptoethanol (Gibco), and 10 ng/mL mouse leukemia inhibitory factor (mLIF) on gelatin-coated dishes in the presence of 5% CO_2_ atmosphere at 37°C.

To generate pluripotency EGFP reporter mouse E14 WT and Mcm2-2A ES cell line, the lentivirus-based EGFP reporter vector PL-SIN-EOS-C(3+)-EGFP plasmid (21318, Addgene) was used for infection. After selection, single cells were seeded and individual clones were then isolated, expanded and confirmed under fluorescence microscope.

### Antibodies

Antibodies used in this study were: anti-MERVL-gag (A-2801, Epigentek), anti-Oct4 (sc-5279; Santa Cruz Biotechnology), anti-Nanog (A300-397A, Bethyl), anti-Tubulin (12G10, DSHB), anti-Ezh2 (5246, Cell Signaling), anti-H3K4me3 (ab8580, Abcam), anti-H3K27me3 (9733, Cell Signaling), anti-Flag (F1804, Sigma Aldrich), anti-Mcm2 (ab4461, Abcam) and anti-Flag (F1804, Sigma).

### Embryoid body (EB) assay

Mouse ES cells were disaggregated and suspended in ES cell medium without LIF. EBs were formed using the hanging drop method (300 cells per drop) on dish lids for 3 days. EBs were then collected and cultured in 10 cm low attachment Petri dish in ES cell culture medium without LIF, and the medium was changed every other day. Samples were collected at the indicated time points for analysis of gene expression.

### Immunofluorescence

Cells were seeded on coverslip coated with 1% gelatin, and then fixed in 4% of formaldehyde for 15 min at room temperature (RT). After washing with PBS, fixed cells were permeabilized with 0.1% Triton X-100 in PBS (PBST) for 10 min and blocked for 1 hour with 5% normal goat serum (NGS) in PBST at RT. Cells were incubated with primary antibodies diluted in 1% NGS in PBST overnight at 4°C. Cells were then washed with PBST and incubated with fluorophore-labeled secondary antibodies for 1 hour at RT. DNA were stained with DAPI. Images were captured by Nikon 80i Fluorescence Microscope.

### RT-PCR analysis

Total RNA was isolated from 1 × 10^6^ cells using RNeasy Plus kit (74104, Qiagen). 0.5 μg of total RNA were used for cDNA synthesis with random hexamers (18080-051, Invitrogen). Quantitative PCR was performed in 12 μL reactions containing 0.1 μM primers and SYBR Green PCR Master Mix (Bio-Rad Laboratories). Primers used are listed below.

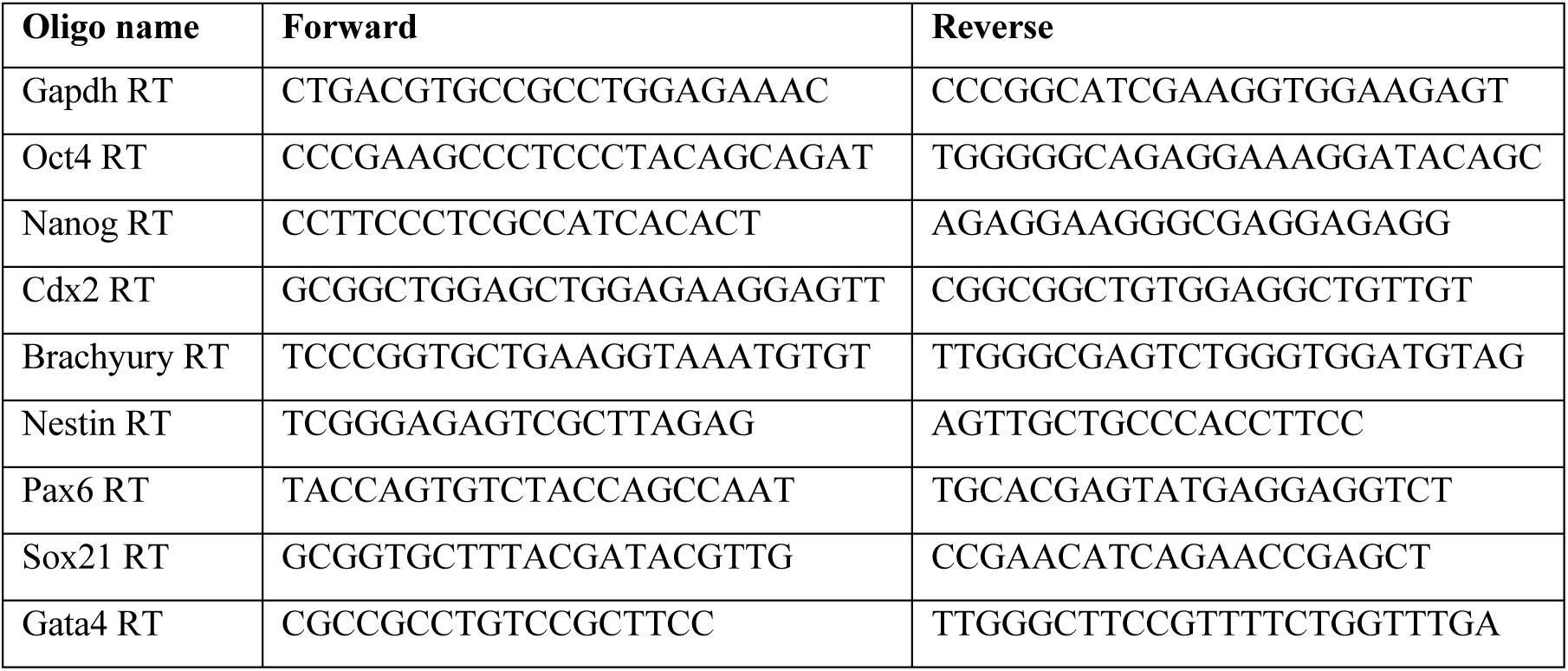

### Neural differentiation assay

The neural differentiation of ES cells was performed as previously described with some modifications (Gao *et al*., 2018; Ying et al., 2003). Briefly, ES cells (day 1) were seeded at high density (2 ×10^6^ cells/60 mm dish) onto gelatin-coated dish in standard mES medium with LIF for 24 hours. The differentiation was initiated by seeding ES cells (day 2) in N2B27 medium at a density of 3 ×10^5^ cells per 60 mm dish. The N2B27 medium was changed every day up to day 6. Cells were cultured continuously in N2B27 medium supplied with EGF (10 ng/ml, R&D) and FGF-2 (10 ng/ml, R&D) for three more days with medium changed every day. Neural precursors were collected at day 9 for analysis.

### FACS analysis of GFP ratio and BrdU incorporation

For GFP ratio analysis, exponentially growing mouse ES cells expressing EGFP reporter gene driven by Oct4 distal enhancer were collected and washed in PBS. Samples were analyzed by Attune NxT software of Attune flow cytometer (Thermo Fisher Scientific). Data were analyzed by FCS Express (version 7).

### CUT&RUN

CUT&RUN was performed by following a published protocol with some modifications (Skene et al., 2018). Cells were fixed in 0.5% PFA for 2 mins and washed three times with washing buffer (20 mM HEPES-NaOH pH 7.5, 150 mM NaCl, 1% Triton X-100, 0.05% SDS, 0.5 mM Spermidine and 1× proteinase inhibitor cocktail) and immobilized to concanavalin A–coated magnetic beads. Cells were then incubated overnight at 4°C with primary antibody (1:400 for Mcm2, 1:1000 for Flag, 1:1000 for H3K4me3 and 1:400 for H3K27me3) in antibody binding buffer (20 mM HEPES-NaOH pH=7.5, 150 mM NaCl, 0.5 mM Spermidine, 1% Triton X-100, 0.05% SDS, 2 mM EDTA, 0.04% Digitonin and 1× proteinase inhibitor cocktail). After washing with dig-wash buffer (0.04% Digitonin in washing buffer), cells were incubated with pre-assembled 2^nd^ antibody+pA-MNase complex in dig-wash buffer for 1 hour at 4°C. After washing unbound pA-MNase, 2 mM CaCl_2_ was added to initiate digestion at 0°C for 30 mins. Reactions were stopped by mixing with 2XStop buffer (340 mM NaCl, 20 mM EDTA, 4 mM EGTA, 0.05% Digitonin, 25 μL 100 µg/ml RNase A, 1% Triton X100 and 0.05% SDS) followed by incubation at 37℃ for 30 mins. DNA in supernatant was mixed with same volume of 2X Elution buffer (20 mM Tris-HCl pH=8.0, 20 mM EDTA, 300 mM NaCl, 10 mM DTT and 2% SDS) and reverse crosslinked at 65℃ overnight. DNA was purified using the QIAquick PCR Purification Kit (28104, Qiagen) and used for library preparation with the Accel-NGS 1S Plus DNA library kit (Swift Bioscience, 10096). Each of the library DNAs were sequenced using paired-end sequencing by Illumina NextSeq 500 platforms at the Columbia University Genome Center.

### ATAC-Seq

Cells were collected, washed once with cold PBS, and lysed in cell lysis buffer (10 mM Tris-HCl pH 7.5, 10 mM NaCl, 3 mM MgCl_2_, 0.1% NP-40, 0.1% Tween-20, 0.01% Digitonin). After incubating on ice for 3 mins, cells were washed once in wash buffer (10 mM Tris-HCl pH 7.5, 10 mM NaCl, 3 mM MgCl_2_, 0.1% Tween-20), and centrifuge at 500 g for 10 mins. Cell pellets were then incubated in transposition reaction buffer (10 mM Tris-HCl pH 7.6, 5 mM MgCl_2_, 10% Dimethylformamide, 0.1% Tween-20, 0.01% Digitonin, 33% 1XPBS) with 1.5 ul pAG-Tn5 (15-1017, EpiCypher) at 37°C for 30 mins. DNA was purified using the QIAquick PCR Purification Kit (28104, Qiagen). Library PCR was performed using standard Illumina Nextera Dual Indexing primers. Libraries were sequenced using paired-end sequencing on Illumina NextSeq 500 platforms at the Columbia University Genome Center.

### CUT&RUN and ATAC-Seq analysis

CUT&RUN and ATAC libraries were constructed and sequenced using paired-end method on Illumina platforms. Adaptor sequences of all raw reads were removed by Cutadapt (Marcel, 2011) and reads <10nt were removed. CUT&RUN and ATAC-Seq data were then mapped to mouse (mm10) reference genome by Bowtie2 (Langmead and Salzberg, 2012). Multi-mapped reads were removed using SAMtools (MAPQ<40] (Li et al., 2009) and duplicate reads were removed using Sambamba software (Tarasov et al., 2015). Read coverage in a bin of 1bp was calculated from filtered bam files by deepTools2 (Ramirez et al., 2016) and then normalized with total filtered reads number into reads per million (RPM). Genome wide correlation was performed by deepTools2 (Ramirez *et al*., 2016) in a bin of 5000bp. Peaks were called by MACS (Zhang et al., 2008) by parameters “macs2 callpeak -g mm -f BAMPE -p 1e-04 --broad --broad-cutoff 1e-04 --llocal 10000 – nolambda” and the cutoff of peak was p=0.0001.

Read density level surrounding gene promoters ([-3kb, 3kb] of TSS) or MCM2 peaks was calculated by featureCounts (Liao et al., 2014) and then normalized to reads per kilobase per million reads (RPKM). Heatmap clustering was performed by “ward.D2” (Legendre, 2014) method based on z score of log10(RPKM). To identify differential CUT&RUN peaks, peaks from both wild type and mutant cells were firstly merged to a union pool and then read counts were calculated in the merged peaks by featureCounts (Liao *et al*., 2014). DESeq2 (Love et al., 2014) was then used to identify differential peaks by |log2 fold change|>1.

### RNA-seq analysis

Total RNA was isolated from 1 × 10^6^ cells using RNeasy Plus Mini kit (74136, Qiagen). RNA-seq libraries were prepared and deep sequenced in Columbia University Genome Center. Two replicates for each sample were sequenced. RNA-seq libraries were sequenced using paired-end method on Illumina platforms. Adaptor sequences of all raw reads were removed by Cutadapt (Marcel, 2011) and reads <10nt were removed. RNA-seq data were mapped to the mouse (mm10) reference genome by STAR software (Dobin et al., 2013). Gene expression levels were firstly calculated by featureCounts (Liao *et al*., 2014) to obtain read counts and then normalized with total mapped reads into reads per kilobase in per million reads (RPKM). Differential expressed genes were identified by DESeq2 (Love *et al*., 2014) by using adjusted p value<0.01 and |log2 fold change|>1. Gene ontology enrichment analysis was performed by “cluserProfiler” (Yu et al., 2012) in a level 4 of “biological process” functions.

### Statistical analyses

Data are presented as means ± SD. Differences between groups were evaluated using two-tailed unpaired Student t test (noted in figure legends). Statistical analysis was performed in GraphPad Prism software (version 7). All tests were considered significant at p < 0.05. For all the sequenced data analysis, the statistical test was performed using R software. Difference test between groups were evaluated by Wilcoxon signed-rank test.

## DATA AND SOFTWARE AVAILABILITY

Raw and processed sequencing data generated in the course of this study can be accessed via the GEO database with accession number: GSE203272. The token for reviewer access is : cxerwmoslrkflmt.

## Acknowledgements

We thank Richard He for critical editing of the paper. DNA sequencing was performed in Columbia Genome Center with the support from the Herbert Irving Comprehensive Cancer Center at Columbia University and supported by NIH/NCI Cancer Center Support Grant P30CA013696. This work is supported by NIH grant GM R35118015 (to Z.Z.).

## Author Contributions

X.X. and Z.Z. conceived the project. X.X. designed experiments. X.X., K.B. and X.R. performed experiments. X.H. performed the data analysis. Z.Z. supervised the study. X.X., X.H. and Z.Z. wrote the manuscript with comments from all authors.

## Supplementary Figure

**Figure 1-figure supplement 1:**
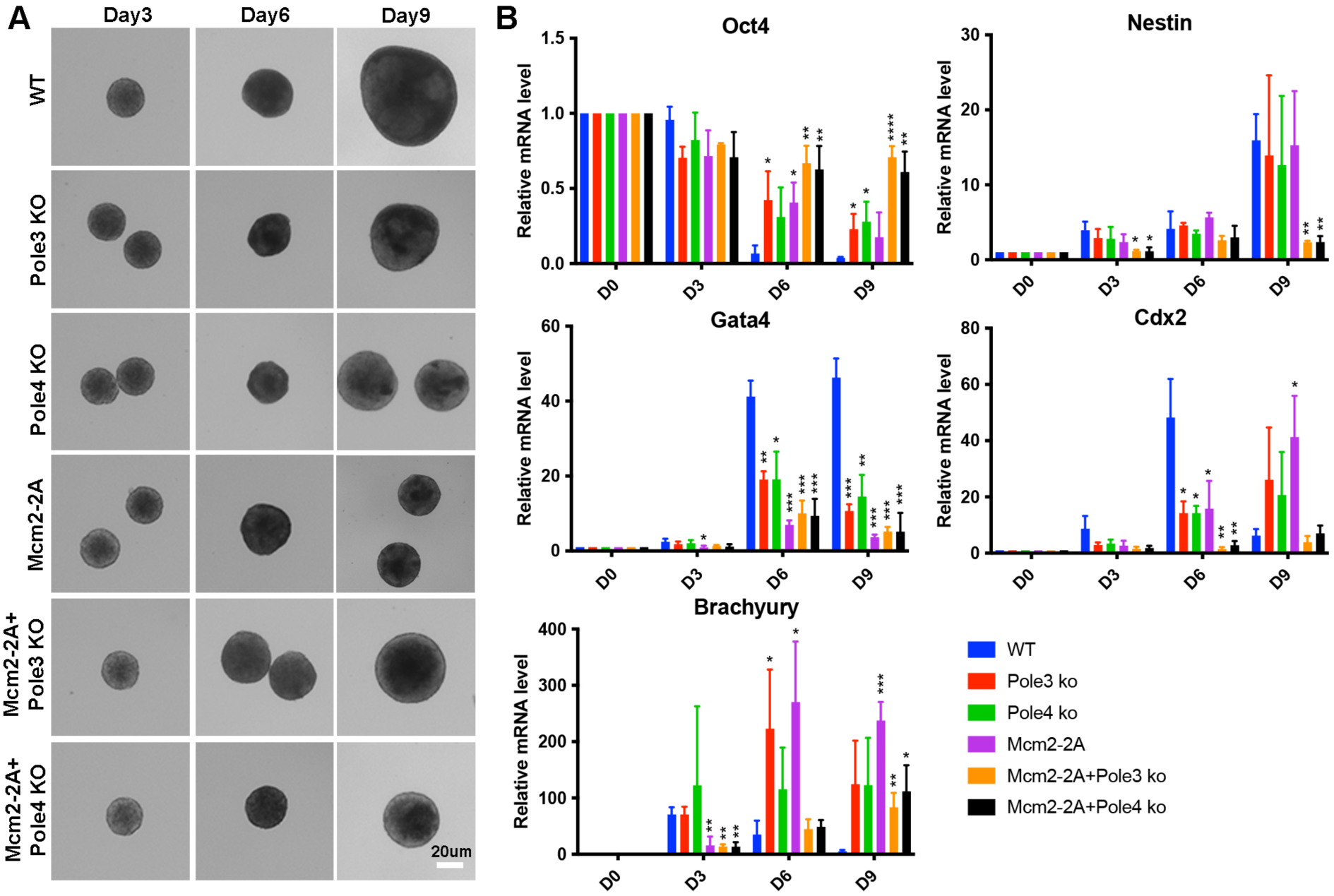
Mcm2-2A, Pole3 KO and Pole4 KO mutations affect ES cell differentiation based on in vitro embryonic body (EB) formation assays. See also Figure 1-figure supplement 1-source data 1. (A) Representative images of WT, Pole3 KO, Pole4 KO, Mcm2-2A, Mcm2-2A + Pole3 KO and Mcm2-2A + Pole4 KO cells during the process of EB formation. Scale bar: 20 µm. (B) RT-PCR analysis of expression of *Oct4* (a gene involved in pluripotency) and four lineage-specific genes (*Nestin*, *Gata4*, *Cdx2* and *Brachyury*) in WT, Pole3 KO, Pole4 KO, Mcm2-2A, Mcm2-2A+Pole3 KO and Mcm2-2A+Pole4 KO cells during EB formation. The relative expression of each gene against house-keeping gene GAPDH is presented as means ± SD from three independent experiments. Statistical analysis was performed by two-tailed unpaired Student *t* test (\**P* < 0.05; \*\**P* < 0.01; \*\*\**P* < 0.001).

**Figure 2-figure supplement 1:**
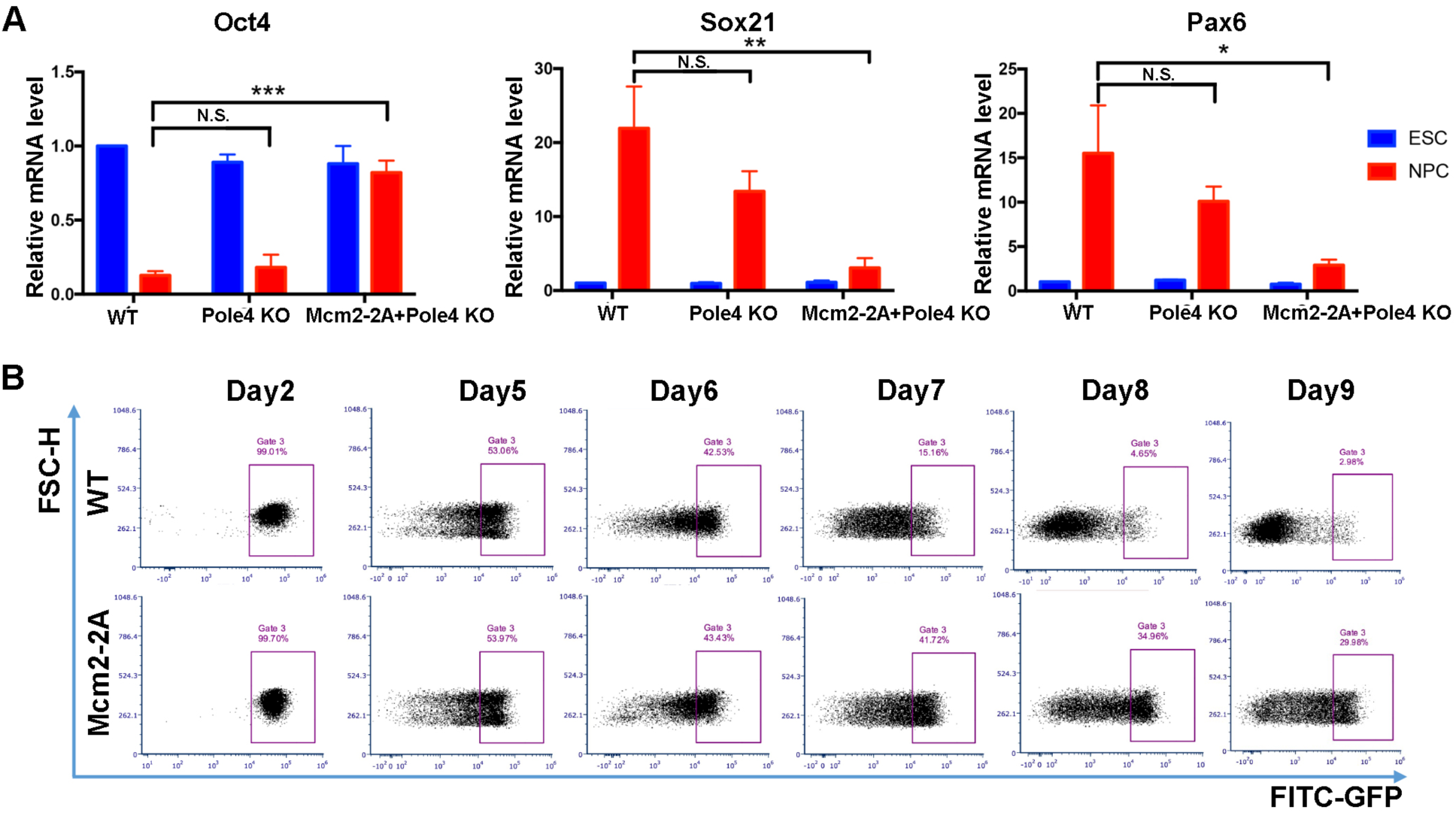
Mcm2-2A mutant prohibits neural differentiation of mouse ES cells. See also Figure 2-figure supplement 1-source data 1. (A) RT-PCR analysis of expression of one pluripotency gene (*Oct4*) and two neural lineage specific genes (*Pax6*, *Sox21*) in WT, Pole4 KO and Mcm2-2A + Pole4 KO cells during neural differentiation. GAPDH was used for normalization. Data are presented as means ± SD from three independent experiments (\**P* < 0.05; \*\**P* < 0.01; \*\*\**P* < 0.001; N.S., no significant difference). (B) FACS analysis of the percentage of GFP+ cells in WT and Mcm2-2A cells during neural differentiation. The expression of EGFP is driven by the Oct4 distal enhancer.

**Figure 4-figure supplement 1:**
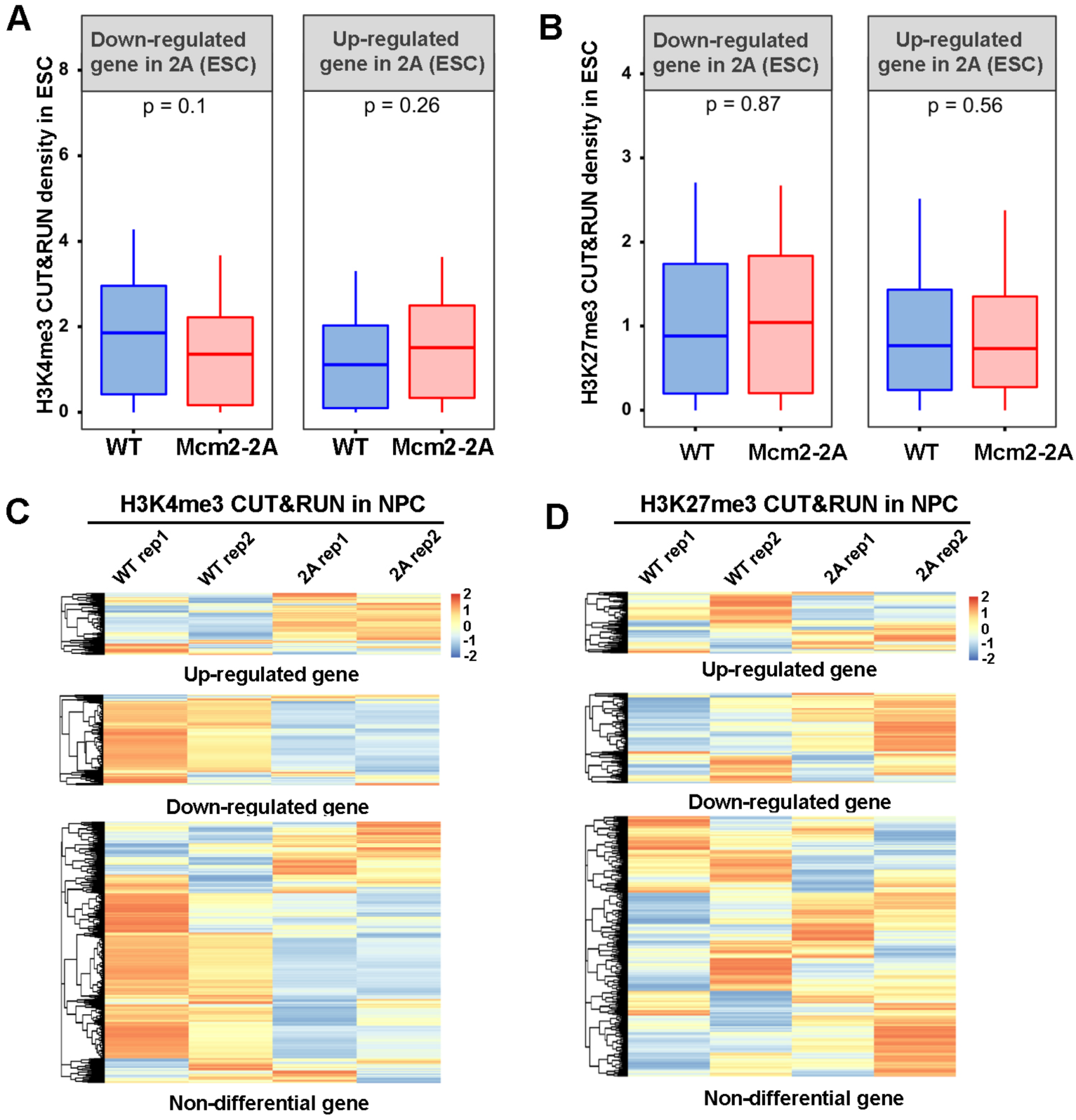
Effects of Mcm2-2A mutation on H3K4me3 and H3K27me3 distribution during neural differentiation. (A and B) Average of H3K4me3 (A) and H3K27me3 (B) CUT&RUN density in WT and Mcm2-2A ES cells at the promoters ([-3k, 3k] around TSS) of down-regulated and up-regulated genes in Mcm2-2A ES cells compared to wild type ES cells (Figure 3A, left). The Y-axis represents the log2 ratio of CUT&RUN density (RPKM). The *P* values were calculated using Wilcoxon signed-rank test from two independent replicates. (C and D) Heatmap of H3K4me3 (C) and H3K27me3 (D) CUT&RUN density at the promoters ([-3k, 3k] around TSS) of up-regulated, down-regulated and non-differential genes in Mcm2-2A NPCs (Figure 3A, right). Two independent replicates were shown for each cell line.

**Figure 5-figure supplement 1:**
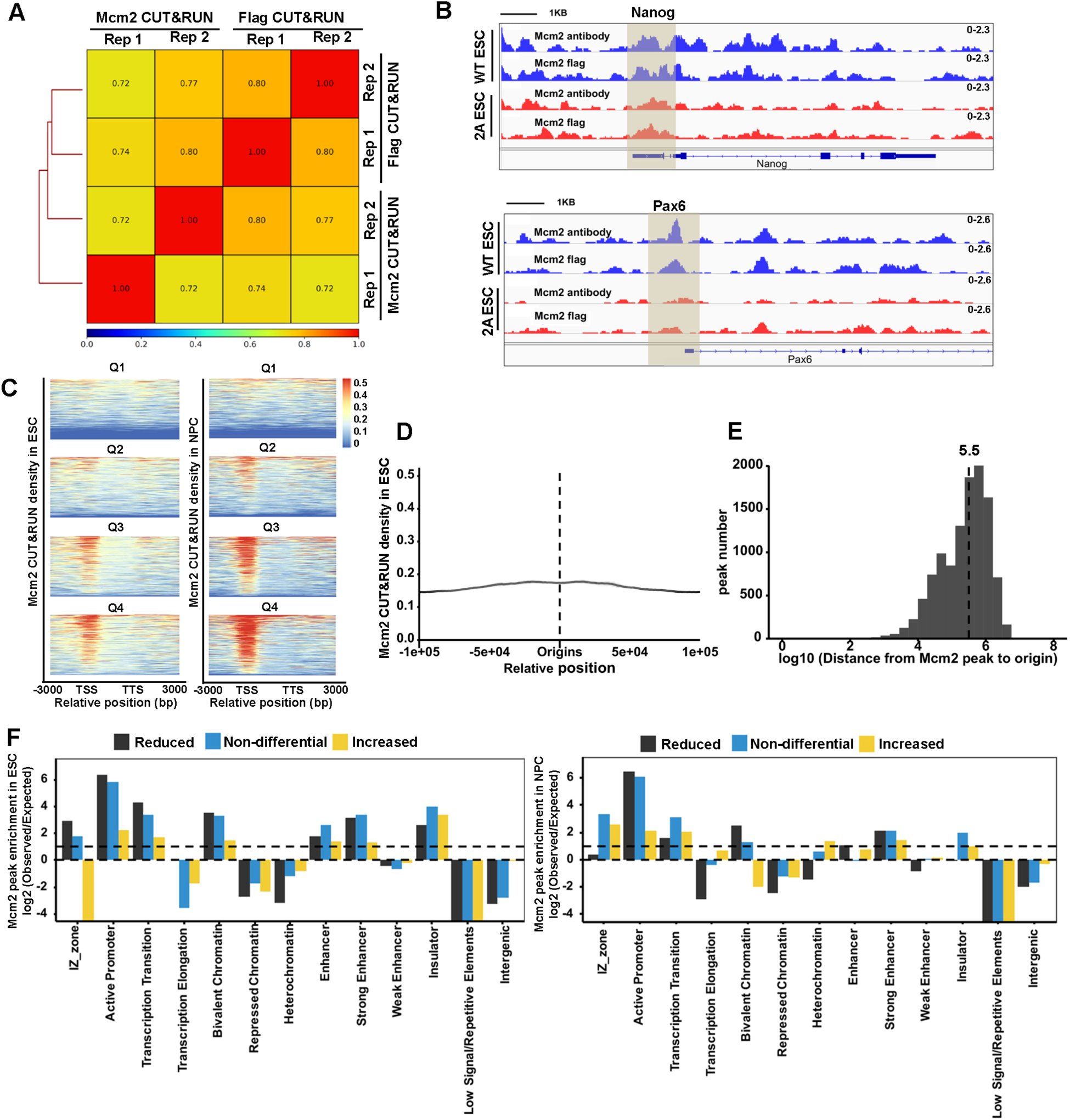
Mcm2-2A mutation decreases Mcm2 binding at chromatin. (A) Genome wide correlations of Mcm2 CUT&RUN datasets generated using antibodies against the Flag epitope fused to Mcm2 or Mcm2-2A or antibodies against Mcm2, each with two independent repeats using a 5 kb sliding window. (B) Snapshots displaying Flag-Mcm2 CUT&RUN and Mcm2 CUT&RUN density at Oct4 and Pax6 loci of Flag-Mcm2 tagged WT and Mcm2-2A ES cells. One representative result from two independent replicates was shown. Shadows indicate CUT&RUN signals around the TSS. (C) Representative heatmaps of Mcm2 CUT&RUN (RPM, reads per million) density surrounding TSS and TTS from highly expressed to lowly expressed genes in WT and Mcm2-2A ES cells (left) and NPCs (right). Genes were separated into 4 groups based on their expression in mouse ES cells (Q1=lowest quartile, Q4=highest quartile). (D) Average density of Mcm2 CUT&RUN (RPM) in WT ES cells surrounding replication origins (N=1548) in mouse ES cells from two independent replicates. (E) The distribution of Mcm2 CUT&RUN peaks based on the distance between Mcm2 peaks to the nearest origins in WT ES cells. The Y-axis represents the number of Mcm2 CUT&RUN peaks. The medium distance of Mcm2 peaks to the nearest origins is ∼316 kb. The average of two independent replicates were shown. (F) Relative enrichment of increased, reduced and non-differential peaks of Mcm2-2A in ES cells (left) and NPCs (right) at different genomic annotations. The Y-axis represents the log2 ratio of the Mcm2 enrichment after normalized to expected ratio of each genomic annotation. Reduced peaks are enriched at active promoters, strong enhancers and bivalent chromatin domains.

**Figure 6-figure supplement 1:**
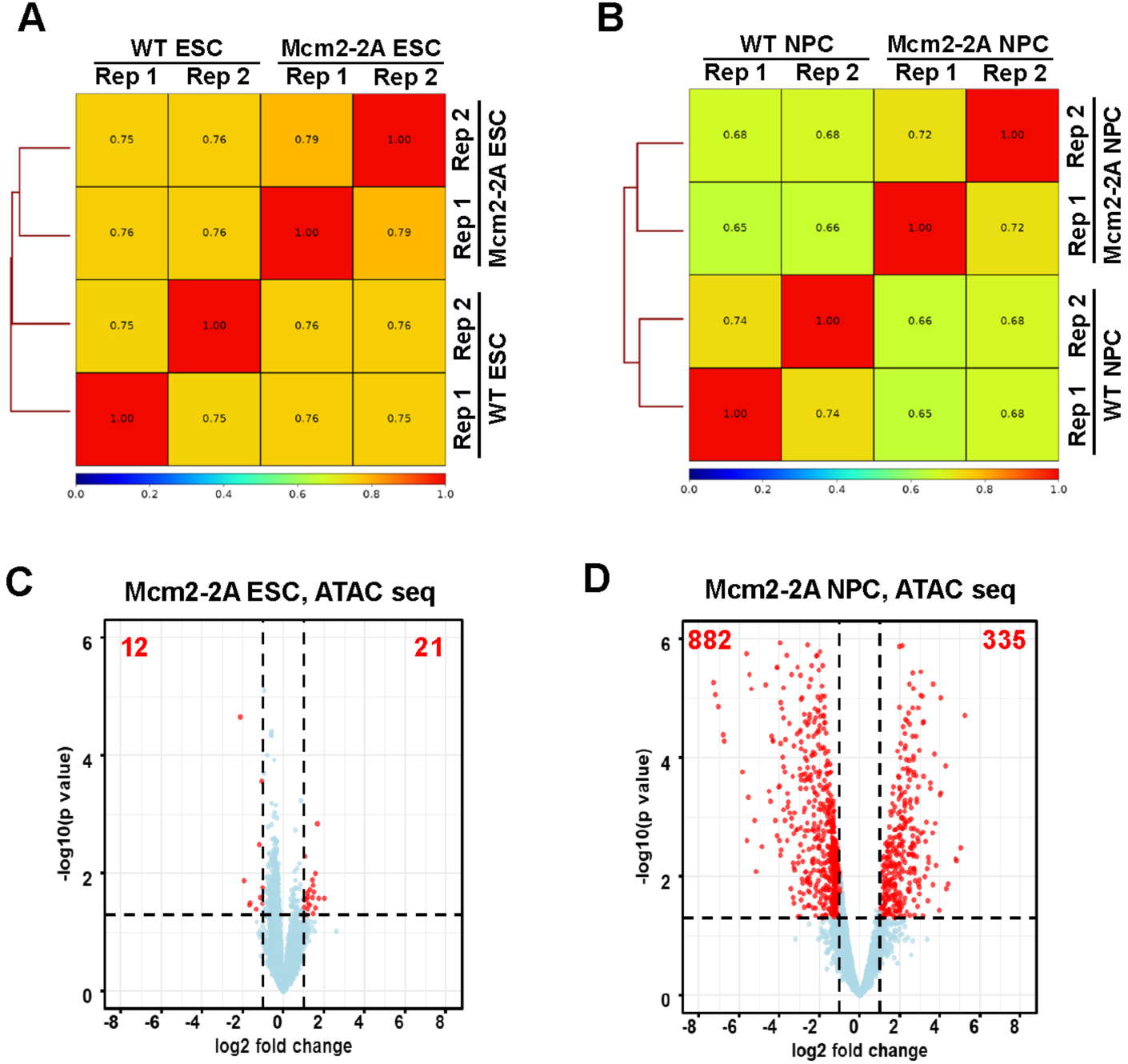
Chromatin accessibility landscape is altered in Mcm2-2A NPCs compared to wild type NPCs. (A and B) Genome wide correlation of ATAC-seq density in WT and Mcm2-2A mutant ES cells (A) and NPCs (B) using a 5 kb window. Two independent replicates were shown for each cell line. (C and D) Volcano plot of differential ATAC-seq peaks between WT and Mcm2-2A mutant ES cells (C) and NPCs (D), with the number of significantly up-regulated and down-regulated peaks (*P*<0.05, |log2 fold change|>1) from two independent replicates shown.

**Figure 6-figure supplement 2:**
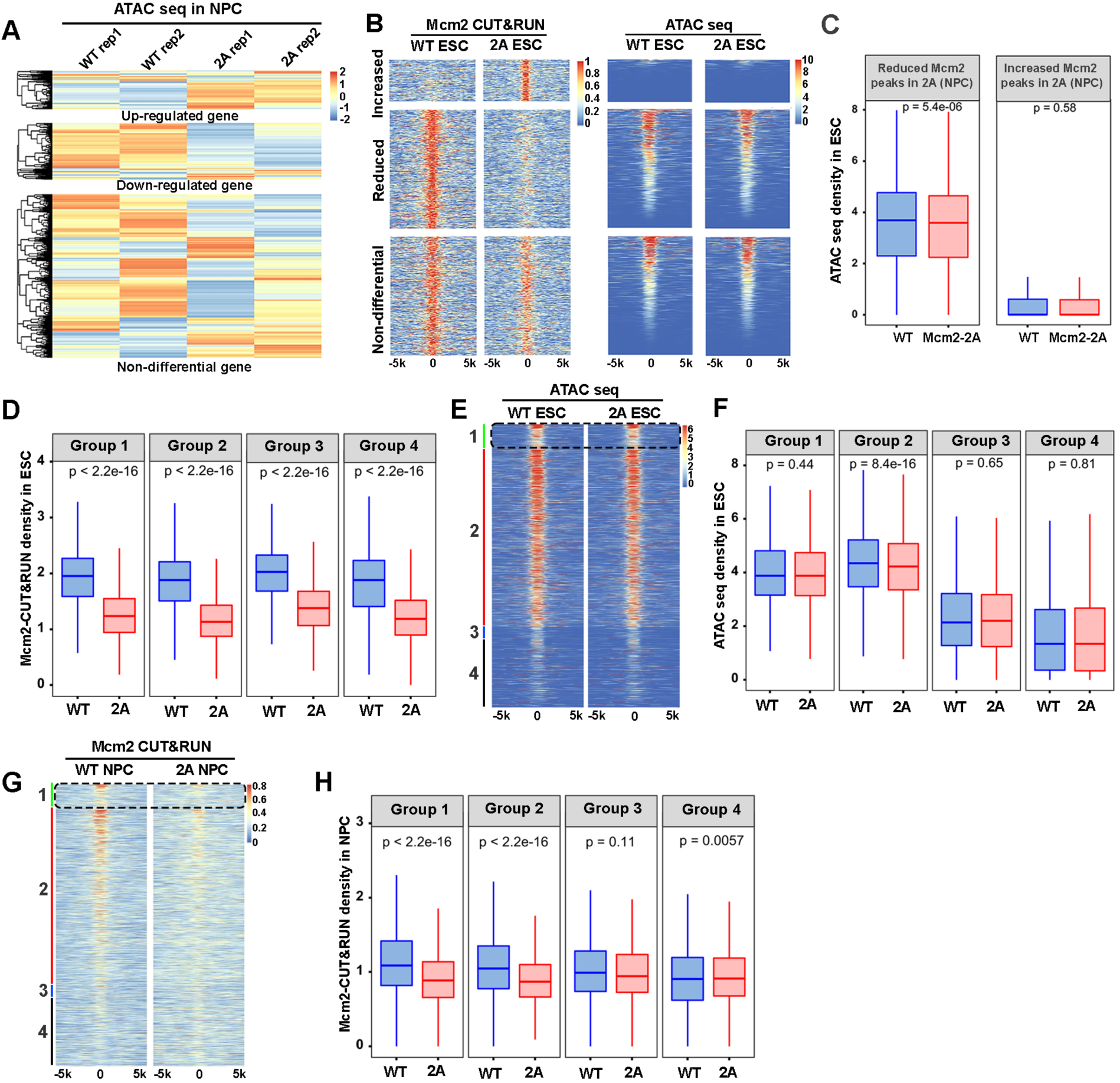
Relationships between the impact of Mcm2-2A mutation on Mcm2 binding and chromatin accessibility. (A) Heatmap of ATAC-seq density at the promoters ([-3k, 3k] around TSS) of up-regulated, down-regulated and non-differential genes in Mcm2-2A NPCs (Figure 3A, right). Two independent replicates were shown for each cell line. (B) Representative heatmap of ATAC-seq density (left) surrounding Mcm2 CUT&RUN peaks (right, |log2 fold change|>1) in WT and Mcm2-2A ES cells. One representative result from two independent replicates was shown. (C) Average ATAC-seq density in WT and Mcm2-2A ES cells at the reduced and increased Mcm2 CUT&RUN peaks in Mcm2-2A ES cells shown in B. (D) Average Mcm2 CUT&RUN density in WT and Mcm2-2A ES cells at each of the four groups of Mcm2 peaks classified in ES cells shown in main Figure 5A. (E) Representative heatmap of ATAC-seq peaks in WT and Mcm2-2A ES cells at four groups of Mcm2 peaks classified in Figure 5A. (F) Average ATAC-seq density in WT and Mcm2-2A ES cells at each of the four groups of Mcm2 peaks classified in Figure 5A. (G) Representative heatmap of Mcm2 CUT&RUN peaks in wild type and Mcm2-2A NPCs at four groups of Mcm2 peaks classified in Figure 5A. (H) Average Mcm2 CUT&RUN density in WT and Mcm2-2A NPCs at each of the four groups of Mcm2 peaks classified in Figure 5A. (C, D, F and H) The Y-axis represents the log2 ratio of ATAC-seq density (C and F, RPKM) or Mcm2 CUT&RUN density (D and H, RPKM). The *P* values were calculated using Wilcoxon signed-rank test. The average of two independent replicates is shown.

## Source data files

**Figure 1-source data 1.** Relative mRNA level of pluripotency and lineage specific genes during EB formation in WT and Mcm2-2A cells, and 2C-like ratio in WT and Mcm2-2A ESCs.

**Figure 1-figure supplement 1-source data 1.** Relative mRNA level of pluripotency and lineage specific genes during EB formation in WT, Pole4 KO, Pole4 KO, Mcm2-2A, Mcm2-2A+Pole3 KO and Mcm2-2A+Pole4 KO cells.

**Figure 2-source data 1.** Relative mRNA level of pluripotency and lineage specific genes, and Oct4 enhancer driven EGFP positive cells ratio during neural differentiation in WT and Mcm2-2A cells.

**Figure 2-source data 2.** Whole SDS-PAGE images and uncropped blots represented in Figure 2D. Oct4 and Nanog protein levels in WT and Mcm2-2A ESCs and NPCs.

**Figure 2-figure supplement 1-source data 1.** Relative mRNA level of pluripotency and lineage specific genes during neural differentiation in WT, Pole4 KO and Mcm2-2A+Pole4 KO cells.

**Figure 4-source data 1.** Whole SDS-PAGE images and uncropped blots represented in Figure 4A. H3K4me3, H3K27me3, Ezh2, Mcm2, Oct4 and Nanog protein levels in WT and Mcm2-2A ESCs and NPCs.

## References

Atlasi, Y., and Stunnenberg, H.G. (2017). The interplay of epigenetic marks during stem cell differentiation and development. Nat Rev Genet 18, 643–658. 10.1038/nrg.2017.57.

Banaszynski, L.A., Wen, D.C., Dewell, S., Whitcomb, S.J., Lin, M.Y., Diaz, N., Elsasser, S.J., Chapgier, A., Goldberg, A.D., Canaani, E., et al. (2013). Hira-Dependent Histone H3.3 Deposition Facilitates PRC2 Recruitment at Developmental Loci in ES Cells. Cell 155, 107–120. 10.1016/j.cell.2013.08.061.

Bernstein, B.E., Mikkelsen, T.S., Xie, X., Kamal, M., Huebert, D.J., Cuff, J., Fry, B., Meissner, A., Wernig, M., Plath, K., et al. (2006). A bivalent chromatin structure marks key developmental genes in embryonic stem cells. Cell 125, 315–326. 10.1016/j.cell.2006.02.041.

Cheloufi, S., Elling, U., Hopfgartner, B., Jung, Y.L., Murn, J., Ninova, M., Hubmann, M., Badeaux, A.I., Ang, C.E., Tenen, D., et al. (2015). The histone chaperone CAF-1 safeguards somatic cell identity. Nature 528, 218–224. 10.1038/nature15749.

Cheng, L., Zhang, X., Wang, Y., Gan, H., Xu, X., Lv, X., Hua, X., Que, J., Ordog, T., and Zhang, Z. (2019). Chromatin Assembly Factor 1 (CAF-1) facilitates the establishment of facultative heterochromatin during pluripotency exit. Nucleic Acids Res 47, 11114–11131. 10.1093/nar/gkz858.

Corpet, A., and Almouzni, G. (2009). Making copies of chromatin: the challenge of nucleosomal organization and epigenetic information. Trends Cell Biol 19, 29–41. 10.1016/j.tcb.2008.10.002.

De Koning, L., Corpet, A., Haber, J.E., and Almouzni, G. (2007). Histone chaperones: an escort network regulating histone traffic. Nat Struct Mol Biol 14, 997–1007. 10.1038/nsmb1318.

De Los Angeles, A., Ferrari, F., Xi, R., Fujiwara, Y., Benvenisty, N., Deng, H., Hochedlinger, K., Jaenisch, R., Lee, S., Leitch, H.G., et al. (2015). Hallmarks of pluripotency. Nature 525, 469–478. 10.1038/nature15515.

Dobin, A., Davis, C.A., Schlesinger, F., Drenkow, J., Zaleski, C., Jha, S., Batut, P., Chaisson, M., and Gingeras, T.R. (2013). STAR: ultrafast universal RNA-seq aligner. Bioinformatics 29, 15–21. 10.1093/bioinformatics/bts635.

Edwards, M.C., Tutter, A.V., Cvetic, C., Gilbert, C.H., Prokhorova, T.A., and Walter, J.C. (2002). MCM2-7 complexes bind chromatin in a distributed pattern surrounding the origin recognition complex in Xenopus egg extracts. J Biol Chem 277, 33049–33057. 10.1074/jbc.M204438200.

English, C.M., Adkins, M.W., Carson, J.J., Churchill, M.E., and Tyler, J.K. (2006). Structural basis for the histone chaperone activity of Asf1. Cell 127, 495–508. 10.1016/j.cell.2006.08.047.

Evrin, C., Serra-Cardona, A., Duan, S., Mukherjee, P.P., Zhang, Z., and Labib, K.P.M. (2022). Spt5 histone binding activity preserves chromatin during transcription by RNA polymerase II. EMBO J 41, e109783. 10.15252/embj.2021109783.

Fang, H.T., El Farran, C.A., Xing, Q.R., Zhang, L.F., Li, H., Lim, B., and Loh, Y.H. (2018). Global H3.3 dynamic deposition defines its bimodal role in cell fate transition. Nat Commun 9. ARTN 1537 10.1038/s41467-018-03904-7.

Gan, H., Serra-Cardona, A., Hua, X., Zhou, H., Labib, K., Yu, C., and Zhang, Z. (2018). The Mcm2-Ctf4-Polalpha Axis Facilitates Parental Histone H3-H4 Transfer to Lagging Strands. Mol Cell 72, 140–151 e143. 10.1016/j.molcel.2018.09.001.

Gao, Y., Gan, H., Lou, Z., and Zhang, Z. (2018). Asf1a resolves bivalent chromatin domains for the induction of lineage-specific genes during mouse embryonic stem cell differentiation. Proc Natl Acad Sci U S A 115, E6162–E6171. 10.1073/pnas.1801909115.

Groth, A., Corpet, A., Cook, A.J.L., Roche, D., Bartek, J., Lukas, J., and Almouzni, G. (2007). Regulation of replication fork progression through histone supply and demand. Science 318, 1928–1931. 10.1126/science.1148992.

Harikumar, A., and Meshorer, E. (2015). Chromatin remodeling and bivalent histone modifications in embryonic stem cells. EMBO Rep 16, 1609–1619. 10.15252/embr.201541011.

Hotta, A., Cheung, A.Y.L., Farra, N., Vijayaragavan, K., Seguin, C.A., Draper, J.S., Pasceri, P., Maksakova, I.A., Mager, D.L., Rossant, J., et al. (2009). Isolation of human iPS cells using EOS lentiviral vectors to select for pluripotency. Nat Methods 6, 370–U378. 10.1038/Nmeth.1325.

Huang, H.D., Stromme, C.B., Saredi, G., Hodl, M., Strandsby, A., Gonzalez-Aguilera, C., Chen, S., Groth, A., and Patel, D.J. (2015). A unique binding mode enables MCM2 to chaperone histones H3-H4 at replication forks. Nature Structural & Molecular Biology 22, 618–626. 10.1038/nsmb.3055.

Ishiuchi, T., Enriquez-Gasca, R., Mizutani, E., Boskovic, A., Ziegler-Birling, C., Rodriguez-Terrones, D., Wakayama, T., Vaquerizas, J.M., and Torres-Padilla, M.E. (2015). Early embryonic-like cells are induced by downregulating replication-dependent chromatin assembly. Nat Struct Mol Biol 22, 662–671. 10.1038/nsmb.3066.

Keller, G. (2005). Embryonic stem cell differentiation: emergence of a new era in biology and medicine. Genes Dev 19, 1129–1155. 10.1101/gad.1303605.

Langmead, B., and Salzberg, S.L. (2012). Fast gapped-read alignment with Bowtie 2. Nat Methods 9, 357–359. 10.1038/nmeth.1923.

Legendre, P. (2014). Ward’s Hierarchical Agglomerative Clustering Method: Which Algorithms Implement Ward’s Criterion? Journal of Classification 31, 274–295.

Lei, M., Kawasaki, Y., and Tye, B.K. (1996). Physical interactions among Mcm proteins and effects of Mcm dosage on DNA replication in Saccharomyces cerevisiae. Molecular and Cellular Biology 16, 5081–5090.

Li, H., Handsaker, B., Wysoker, A., Fennell, T., Ruan, J., Homer, N., Marth, G., Abecasis, G., Durbin, R., and Proc, G.P.D. (2009). The Sequence Alignment/Map format and SAMtools. Bioinformatics 25, 2078–2079. 10.1093/bioinformatics/btp352.

Li, Z., Hua, X., Serra-Cardona, A., Xu, X., Gan, S., Zhou, H., Yang, W.S., Chen, C.L., Xu, R.M., and Zhang, Z. (2020). DNA polymerase alpha interacts with H3-H4 and facilitates the transfer of parental histones to lagging strands. Sci Adv 6, eabb5820. 10.1126/sciadv.abb5820.

Liao, Y., Smyth, G.K., and Shi, W. (2014). featureCounts: an efficient general purpose program for assigning sequence reads to genomic features. Bioinformatics 30, 923–930. 10.1093/bioinformatics/btt656.

Love, M.I., Huber, W., and Anders, S. (2014). Moderated estimation of fold change and dispersion for RNA-seq data with DESeq2. Genome Biol 15, 550. 10.1186/s13059-014-0550-8.

Marcel, M. (2011). Cutadapt removes adapter sequences from high-throughput sequencing reads. EMBnet.journal 17, 10–12. https://doi.org/10.14806/ej.17.1.200.

Mikkelsen, T.S., Ku, M., Jaffe, D.B., Issac, B., Lieberman, E., Giannoukos, G., Alvarez, P., Brockman, W., Kim, T.K., Koche, R.P., et al. (2007). Genome-wide maps of chromatin state in pluripotent and lineage-committed cells. Nature 448, 553–560. 10.1038/nature06008.

Petryk, N., Dalby, M., Wenger, A., Stromme, C.B., Strandsby, A., Andersson, R., and Groth, A. (2018). MCM2 promotes symmetric inheritance of modified histones during DNA replication. Science 361, 1389–1391. 10.1126/science.aau0294.

Prioleau, M.N., and MacAlpine, D.M. (2016). DNA replication origins-where do we begin? Gene Dev 30, 1683–1697. 10.1101/gad.285114.116.

Ramirez, F., Ryan, D.P., Gruning, B., Bhardwaj, V., Kilpert, F., Richter, A.S., Heyne, S., Dundar, F., and Manke, T. (2016). deepTools2: a next generation web server for deep-sequencing data analysis. Nucleic Acids Res 44, W160–165. 10.1093/nar/gkw257.

Ray-Gallet, D., Quivy, J.P., Scamps, C., Martini, E.M.D., Lipinski, M., and Almouzni, G. (2002). HIRA is critical for a nucleosome assembly pathway independent of DNA synthesis. Mol Cell 9, 1091–1100. Doi 10.1016/S1097-2765(02)00526-9.

Serra-Cardona, A., Duan, S., Yu, C., and Zhang, Z. (2022). H3K4me3 recognition by the COMPASS complex facilitates the restoration of this histone mark following DNA replication. Science Advances 8, eabm6246. doi:10.1126/sciadv.abm6246.

Serra-Cardona, A., and Zhang, Z. (2018). Replication-Coupled Nucleosome Assembly in the Passage of Epigenetic Information and Cell Identity. Trends Biochem Sci 43, 136–148. 10.1016/j.tibs.2017.12.003.

Sha, K., and Boyer, L.A. (2008). The chromatin signature of pluripotent cells. In StemBook. 10.3824/stembook.1.45.1.

Skene, P.J., Henikoff, J.G., and Henikoff, S. (2018). Targeted in situ genome-wide profiling with high efficiency for low cell numbers. Nat Protoc 13, 1006–1019. 10.1038/nprot.2018.015.

Smith, S., and Stillman, B. (1989). Purification and characterization of CAF-I, a human cell factor required for chromatin assembly during DNA replication in vitro. Cell 58, 15–25. 10.1016/0092-8674(89)90398-x.

Snyder, M., Huang, X.Y., and Zhang, J.J. (2009). The Minichromosome Maintenance Proteins 2-7 (MCM2-7) Are Necessary for RNA Polymerase II (Pol II)-mediated Transcription. J Biol Chem 284, 13466–13472. 10.1074/jbc.M809471200.

Tagami, H., Ray-Gallet, D., Almouzni, G., and Nakatani, Y. (2004). Histone H3.1 and H3.3 complexes mediate nucleosome assembly pathways dependent or independent of DNA synthesis. Cell 116, 51–61. 10.1016/s0092-8674(03)01064-x.

Tarasov, A., Vilella, A.J., Cuppen, E., Nijman, I.J., and Prins, P. (2015). Sambamba: fast processing of NGS alignment formats. Bioinformatics 31, 2032–2034. 10.1093/bioinformatics/btv098.

Trevino, A.E., Sinnott-Armstrong, N., Andersen, J., Yoon, S.J., Huber, N., Pritchard, J.K., Chang, H.Y., Greenleaf, W.J., and Pasca, S.P. (2020). Chromatin accessibility dynamics in a model of human forebrain development. Science 367. 10.1126/science.aay1645.

Tye, B.K. (1999). MCM proteins in DNA replication. Annu Rev Biochem 68, 649–686. 10.1146/annurev.biochem.68.1.649.

Voigt, P., Tee, W.W., and Reinberg, D. (2013). A double take on bivalent promoters. Genes Dev 27, 1318–1338. 10.1101/gad.219626.113.

Woodward, A.M., Gohler, T., Luciani, M.G., Oehlmann, M., Ge, X.Q., Gartner, A., Jackson, D.A., and Blow, J.J. (2006). Excess Mcm2-7 license dormant origins of replication that can be used under conditions of replicative stress. J Cell Biol 173, 673–683. DOI 10.1083/jcb.200602108.

Xu, X., Duan, S., Hua, X., Li, Z., He, R., and Zhaang, Z. (2022). Stable inheritance of H3.3-containing nucleosomes during mitotic cell divisions. Nat Commun 13, 2514. 10.1038/s41467-022-30298-4.

Yankulov, K., Todorov, I., Romanowski, P., Licatalosi, D., Cilli, K., McCracken, S., Laskey, R., and Bentley, D.L. (1999). MCM proteins are associated with RNA polymerase II holoenzyme. Mol Cell Biol 19, 6154–6163. 10.1128/MCB.19.9.6154.

Ying, Q.L., Stavridis, M., Griffiths, D., Li, M., and Smith, A. (2003). Conversion of embryonic stem cells into neuroectodermal precursors in adherent monoculture. Nat Biotechnol 21, 183–186. 10.1038/nbt780.

Young, R.A. (2011). Control of the embryonic stem cell state. Cell 144, 940–954. 10.1016/j.cell.2011.01.032.

Yu, C., Gan, H., Serra-Cardona, A., Zhang, L., Gan, S., Sharma, S., Johansson, E., Chabes, A., Xu, R.M., and Zhang, Z. (2018). A mechanism for preventing asymmetric histone segregation onto replicating DNA strands. Science 361, 1386–1389. 10.1126/science.aat8849.

Yu, G., Wang, L.G., Han, Y., and He, Q.Y. (2012). clusterProfiler: an R package for comparing biological themes among gene clusters. OMICS 16, 284–287. 10.1089/omi.2011.0118.

Zhang, Y., Liu, T., Meyer, C.A., Eeckhoute, J., Johnson, D.S., Bernstein, B.E., Nusbaum, C., Myers, R.M., Brown, M., Li, W., and Liu, X.S. (2008). Model-based analysis of ChIP-Seq (MACS). Genome Biol 9, R137. 10.1186/gb-2008-9-9-r137.

